# The role of trehalose 6-phosphate in shoot branching – local and non-local effects on axillary bud outgrowth in arabidopsis rosettes

**DOI:** 10.1101/2020.06.18.158568

**Authors:** Franziska Fichtner, Francois F. Barbier, Maria G. Annunziata, Regina Feil, Justyna J. Olas, Bernd Mueller-Roeber, Mark Stitt, Christine A. Beveridge, John E. Lunn

**Affiliations:** Max Planck Institute of Molecular Plant Physiology, 14476 Potsdam-Golm, Germany; School of Biological Sciences, The University of Queensland, St. Lucia, QLD 4072, Australia; University of Potsdam, Institute of Biochemistry and Biology, Karl-Liebknecht-Straße 24-25, Haus 20, 14476 Potsdam, Germany

**Keywords:** *Arabidopsis thaliana* (arabidopsis), axillary bud, branching, sucrose, sugar signalling, trehalose 6-phosphate

## Abstract

- Trehalose 6-phosphate (Tre6P) is a sucrose signalling metabolite that has been implicated in regulation of shoot branching, but its precise role is not understood.
- We expressed tagged forms of TREHALOSE-6-PHOSPHATE SYNTHASE1 (TPS1) to determine where Tre6P is synthesized in arabidopsis (*Arabidopsis thaliana*), and investigated the impact of localized changes in Tre6P levels, in axillary buds or vascular tissues, on shoot branching in wild-type and branching mutant backgrounds.
- TPS1 is expressed in axillary buds and the subtending vasculature, as well as in the leaf and stem vasculature. Expression of a heterologous trehalose-6-phosphate phosphatase (TPP) to lower Tre6P in axillary buds strongly delayed bud outgrowth in long days and inhibited branching in short days. TPP expression in the vasculature also delayed lateral bud outgrowth and decreased branching. Increased Tre6P in the vasculature enhanced branching and was accompanied by higher expression of *FLOWERING LOCUS T* (*FT)* and up-regulation of sucrose transporters. Increased vascular Tre6P levels enhanced branching in *branched1* but not in *ft* mutant backgrounds.
- These results provide direct genetic evidence of a local role for Tre6P in regulation of axillary bud outgrowth within the buds themselves, and also connect Tre6P with systemic regulation of shoot branching via FT.

## INTRODUCTION

Shoot architecture is a highly plastic trait that influences plant fitness in the wild and the productivity of crop plants (Patrick and Colyvas, 2014). In many plants, the shoot apex inhibits the outgrowth of axillary (lateral) buds, prioritizing allocation of resources towards growth of the main stem. This phenomenon is known as apical dominance. Removal of the shoot apex, by herbivory or pruning, leads to the release of dormancy in axillary buds allowing them to grow out to form new branches. Since the 1930s, there has been a general consensus that auxin is the main signal responsible for apical dominance (Barbier et al., 2019). Auxin is produced in the young leaves at the shoot tip and transported down towards the roots, inhibiting the outgrowth of axillary buds along the shoot (Dun et al., 2009; Muller and Leyser, 2011; Brewer et al., 2013). Although the polar auxin stream in the stem inhibits bud outgrowth, auxin derived from the shoot tip does not itself move into axillary buds. Instead it acts via two other phytohormones: strigolactones and cytokinins (Wickson and Thimann 1958; Sachs and Thimann 1967; Gomez-Roldan et al., 2008; Umehara et al., 2008), which have antagonistic effects on axillary buds (Brewer et al., 2009; Ferguson and Beveridge, 2009; Dun et al., 2012, 2013). Auxin enhances synthesis of strigolactones, which inhibit axillary bud growth (Shimizu-Sato et al., 2009; Domagalska et al., 2011), partly via induction of the BRANCHED1 (BRC1) transcription factor, which is a repressor of branching (Aguilar-Martinez et al., 2007; Braun et al., 2012; Dun et al., 2009, 2012). Conversely, auxin inhibits biosynthesis of cytokinins, which promote axillary bud outgrowth, in part by repression of *BRC1* expression (Braun et al., 2012; Dun et al., 2012). A more recent concept is the auxin canalization model. This postulates that the inability of axillary buds to export auxin produced in the bud is responsible for their dormancy, and that outgrowth occurs when the buds are able to establish their own polar auxin stream (Prusinkiewicz et al., 2009; Muller and Leyser, 2011).

Although these phytohormones undoubtedly play a major role in regulation of shoot branching, decapitation experiments in garden pea (*Pisum sativum*) questioned their involvement in the initial outgrowth of axillary buds. Decapitation leads to changes in the sucrose supply at the level of the lower buds that are highly correlated with the timing of bud outgrowth (Mason et al., 2014; Fichtner et al., 2017). Increasing the sucrose supply to axillary buds by feeding sucrose exogenously or removing the rapidly expanding leaves (i.e. competing sinks) also triggered bud growth, even when the shoot apex itself was left intact to maintain polar auxin flow (Mason et al., 2014). Exogenous sucrose also triggered bud outgrowth in isolated stem sections from pea, rose (*Rosa hybrida*) and arabidopsis (Barbier et al., 2015b; Fichtner et al., 2017), even in the presence of auxin in the growth medium (Bertheloot et al., 2020). Together, these studies provide direct evidence that changes in sucrose supply are the initial signal that releases bud dormancy after decapitation. Observations in other species are consistent with sucrose supply playing an important role in shoot branching (Kebrom et al., 2010; 2012; Barbier et al., 2015a; Martín-Fontecha et al., 2018; Barbier et al., 2019b), with dormant buds displaying a carbon starvation-like transcript profile (Tarancón et al., 2017).

Trehalose 6-phosphate (Tre6P) is an essential signal metabolite in plants. The sucrose-Tre6P nexus model proposes that Tre6P signals the availability of sucrose (Lunn et al., 2006) and acts as a negative feedback regulator of sucrose levels (Yadav et al., 2014; Figueroa and Lunn, 2016). Tre6P is synthesized from UDP-glucose and glucose 6-phosphate by Tre6P synthase (TPS), and dephosphorylated to trehalose by Tre6P phosphatase (TPP). In arabidopsis, transcripts of several *TPS* and *TPP* genes have been detected in meristems and axillary buds, and the expression of many of these genes is influenced by sugars and phytohormones (Osuna et al., 2007; Ramon et al., 2009; Yadav et al., 2014). Transcriptomic and phenotypic analyses of various mutants have implicated Tre6P in shoot and inflorescence branching in arabidopsis (Schluepmann et al., 2003), and in other species (Kebrom and Mullet, 2016). In maize (*Zea mays*), inflorescence branching is increased by mutations in two *TPP* genes – *ZmRAMOSA3* and *ZmTPP4* – that disrupt putative Tre6P signalling functions (Satoh-Nagasawa et al., 2006; Claeys et al., 2019).

We recently demonstrated that Tre6P rapidly accumulates in pea axillary buds after decapitation, and that the rise in bud Tre6P levels following decapitation is dependent on sucrose (Fichtner et al., 2017). Although the rise in bud Tre6P levels coincided with the onset of bud outgrowth, its physiological significance is not yet known (Fichtner et al., 2017). In maize, *grassy tillers1* and *teosinte branched1* mutants are highly branched because the tiller buds fail to establish dormancy, and this trait is associated with tiller buds having elevated levels of Tre6P (Dong et al., 2019).

There is circumstantial evidence that Tre6P might also have a more remote influence on axillary buds and shoot branching, in particular via effects on the expression of the florigenic protein FLOWERING LOCUS T (FT; Wahl et al., 2013). FT is synthesized in the phloem companion cells in leaves and moves in the phloem sieve elements to the shoot apical meristem (SAM), where it interacts with the FLOWERING LOCUS D protein to promote the floral transition (Turck et al., 2008). In arabidopsis, FT and its close homologue, TWIN SISTER OF FT (TSF), interact with the branching repressor BRC1 in axillary buds (Niwa et al., 2013). In rice (*Oryza sativa*), a homologue of FT has been shown to regulate tillering (Tsuji et al., 2015), and there is evidence for an increase in branching mediated by up-regulation of FT in tomato (*Solanum lycopersicum*; Weng et al., 2016) or expression of a heterologous FT in tobacco (*Nicotiana tabacum*; Li et al., 2015). In pea, one FT homologue, GIGAS, has also been implicated in the regulation of bud outgrowth (Beveridge and Murfet, 1996; Hecht et al., 2011).

We hypothesize that Tre6P influences shoot branching in multiple ways, acting both locally within axillary buds and more remotely in the phloem-loading zone of leaves, potentially linking axillary bud outgrowth to the local availability of sucrose as well as the overall C-status of the plant. The aims of this study were to define where Tre6P acts in regulation of axillary bud outgrowth and to identify interactions with other signalling pathways that affect shoot branching. We used tissue-specific promoters to bring about localized changes in Tre6P levels in arabidopsis, and investigated the impact of these changes on shoot branching. We demonstrate that Tre6P plays a central role in the release of axillary bud dormancy to form new shoot branches.

## MATERIALS AND METHODS

### Materials

Wild type arabidopsis (*Arabidopsis thaliana* [L.] Heynh.) accession Columbia-0 (Col-0) seeds were from an in-house collection. The *tps1-1* lines complemented with β-GLUCURONIDASE- or GFP-tagged forms of TPS1 or with the *Escherichia coli* TPS (OtsA) were those described in Fichtner et al. (2020).

### Molecular cloning

For expression of genes of interest under the control of the *BRC1* (*BRANCHED1*, At3g18550) promoter, a 1-kbp genomic region upstream of the start codon of the *BRC1* gene was used. The *GLYCINE-DECARBOXYLASE P-SUBUNIT A* (*GLDPA*) promoter from the C4 plant *Flaveria trinervia* was amplified from the previously used FtGLDPA5’-pBI121 plasmid (see Engelmann et al., 2008), resulting in a 1.5− kbp promoter fragment. All promoter sequences were integrated into pGreenII plasmids (www.pgreen.ac.uk; Hellens et al., 2000) that were equipped with the terminator of the agrobacterial *octopine synthase* gene. The *otsA* gene from *E. coli* (strain K12) was amplified from *E. coli* DNA and the genetic sequences of *CeTPP/GOB-1*, codon-optimized for expression in arabidopsis, was synthesized and cloned by GenScript (www.genscript.com) and sub-cloned into the pGreenII plasmid.

### Arabidopsis transformation

Gene constructs were introduced into arabidopsis Col-0 by *Agrobacterium tumefaciens* mediated (floral dipping (Clough and Bent, 1998). Primary transformants were selected by spraying with 0.05% (v/v) glufosinate. Lines that showed a 3:1 segregation of resistant:susceptible individuals in the T_2_ generation, indicating a single transgenic locus, were chosen for further propagation. Progeny from such lines were screened by glufosinate selection in the T_3_ generation to identify homozygous lines for each transgene by survival assays.

### Plant growth conditions

All arabidopsis seeds were stratified for 3 days at 4°C and grown for 7 days on solid half-strength MS plates (Murashige and Skoog, 1962) and then transferred to a 1:1 mixture of vermiculite and soil (Stender, www.stender.de). Plants were grown in controlled environment chambers fitted with fluorescent lamps (Annunziata et al., 2017), with 8-h, 16-h or 18-h photoperiods, 150 μmol m^−2^ s^−1^ irradiance and day/night temperatures of 22°C/18°C. Unless stated otherwise, plants were harvested 10 h after dawn (ZT10). The age of the plants at harvest is indicated for each individual experiment. Bud enriched material was obtained from rosettes by removing all leaves as well as the hypocotyl, leaving only the inner stem regions and shoot apex. For vascular enriched material, the leaf mid-veins of three plants were dissected and pooled. Dormant axillary buds were collected from 15 plants one week after bolting.

### Phenotyping

Flowering time was determined as total leaf number (rosette + cauline leaves). Rosette and cauline leaves were counted separately, and the rosette leaf number was used to determine RI number per leaf. Shoots were counted either when they had finished flowering and fully senesced or every 2 d after bolting. Every shoot with a size ≥0.5 cm was counted.

### Microscopy

*β-Glucuronidase (GUS) reporter assay*: Plants were placed in GUS staining solution (50 mM sodium-phosphate buffer (pH 7), 5 mM K_3_[Fe(CN)_6_], 5 mM K_4_[Fe(CN)_6_], and 1 mM 5-bromo-4-chloro-3-indolyl-beta-D-glucuronic acid (X-Gluc). After incubation at 37°C in the dark overnight, the tissue was destained by washing several times with 70% (v/v) ethanol. Meristems were harvested, fixed and embedded in wax as described in Olas et al. (2019). Leaves were fixed with Technovit^®^ 7100 (Kulzer, www.kulzer-technik.de) according to the manufacturer’s instructions. Sections (4-μm) were cut using a Leica Rotary Microtome RM2265 (Leica Biosystems, www.leicabiosystems.com) and observed under an Olympus BX-51 Epi-Fluorescence Microscope fitted with a DC View III camera and operated using CellSense software (Olympus, www.olympus-lifescience.com). Non-sectioned plant material was examined using either the microscope described above or a Leica Stereomicroscope MZ12.5 fitted with a DC 420 camera and operated with LAS software (Leica Biosystems). *GFP-reporter lines*: GREEN FLUORESCENT PROTEIN (GFP) expression was detected using a Leica TCS SP8 Spectral Laser Scanning Confocal motorized Microscope operated with LAS X software (Leica Biosystems; www.leica.com). Overlays were done using the image processing package Fiji for ImageJ (https://fiji.sc/).

### Immunoblotting

Expression of heterologous proteins was confirmed by immunoblotting as described previously (Martins et al. 2013). The following primary antibodies were used: (i) rabbit anti-OtsA (Martins et al., 2013; 1:3,000 dilution) or (ii) rabbit anti-CeTPP kindly provided by Dr Carlos Figueroa (MPI-MP, Potsdam-Golm, Germany; 1:4,000 dilution).

### Metabolite analysis

Frozen plant tissue was ground to a fine powder at liquid N2 temperature and water-soluble metabolites were extracted as described in Lunn et al. (2006). Tre6P, other phosphorylated intermediates and organic acids were measured by anion-exchange HPLC coupled to tandem mass spectrometry (Lunn et al., 2006), with modifications as described in Figueroa et al. (2016). Sucrose was measured enzymatically (Stitt et al., 1989).

### Gene expression analysis

RNA was extracted using an RNeasy Plant Mini Kit (Qiagen; www.qiagen.com) following the manufacturer’s instructions. For absolute quantification of transcripts, ArrayControl RNA Spikes (Applied Biosystems; www.thermofisher.com/applied/biosystems) were added before RNA extraction and cDNA synthesis (Flis et al., 2015). Contaminating DNA was removed using a TURBO DNA-free kit, and reverse transcription was performed using a SuperScript IV First-Standard Synthesis System Kit (Invitrogen; www.thermofisher.com/Invitrogen). The PCR mix was prepared using Power SYBR Green PCR Master Mix (Applied Biosystems), and qRT-PCR was performed in a 384-well microplate using an ABI PRISM 7900 HT sequence detection system (Applied Biosystems). Transcript abundance was calculated as described in Flis et al. (2015), using spike numbers 1 to 7. Primers used for spike analysis and qRT-PCR are given in Table S1. Gene expression analysis was also performed as described in Barbier et al. (2019a). cDNA was synthesized by reverse-transcription using an iScript™ cDNA Synthesis Kit (Bio-Rad; www.biorad.com). Quantitative Real-Time PCR was performed using a SensiFAST™ SYBR^®^ No-ROX Kit (Bioline; www.bioline.com). Fluorescence was monitored with a CFX384 thermal cycler (Bio-Rad). Gene expression was calculated using the ΔΔCt method corrected by the primer efficiency, and *TUBULIN3* and *ACTIN* (combination of *ACT2, ACT7* and *ACT8*; Table S1) were used for normalization.

### Statistical analysis

Data plotting and statistical analysis were performed using R Studio Version 1.0.143 (www.rstudio.com) with R version 3.6.2 (https://cran.r-project.org/). Data were analysed using an ANOVA based post hoc comparison of means test using the multiple comparison Fisher’s least significant difference (LSD) test. Figures containing micrographs and other images were compiled using Adobe Illustrator, Microsoft PowerPoint 2010 or ImageJ software (https://imagej.nih.gov/ij/).

## RESULTS

### Localization and function of TPS1 in arabidopsis axillary buds

TPS1 is the predominant enzymatic source of Tre6P in arabidopsis, except in the endosperm of developing seeds (Delorge et al., 2015). To assess the potential for Tre6P synthesis in axillary buds, we complemented the embryo-lethal arabidopsis *tps1-1* null mutant with constructs encoding full-length TPS1 proteins tagged at either the N− or C-terminus with β-GLUCURONIDASE (GUS) or GREEN FLUORESCENT PROTEIN (GFP) (Fig. 1a). Expression of the TPS1 fusion proteins was under the control of the endogenous *TPS1* promoter and other potential regulatory elements from the *TPS1* genomic locus, and the complemented lines showed normal embryonic and post-embryonic growth (Fichtner et al., 2020; Fig. 1a). GUS/GFP-tagged TPS1 was detected in axillary buds (Fig. 1b-e), with strong expression also in the subtending vasculature, but there was little or no expression in leaf primordia or the central meristematic zone of the axillary buds (Fig. 1b-e). This was similar to the expression pattern of TPS1 in the main shoot apex. Before bolting, TPS1 was detected in the flanks and rib zone of the SAM, and in the proto-vasculature subtending the SAM (Fig. 1e), consistent with previous studies of the same reporter lines (Fichtner et al., 2020). Similarly, TPS1 was present in the vasculature subtending the inflorescence SAM as well as in cauline axillary meristems (Fig. 1g).

**Fig 1.**
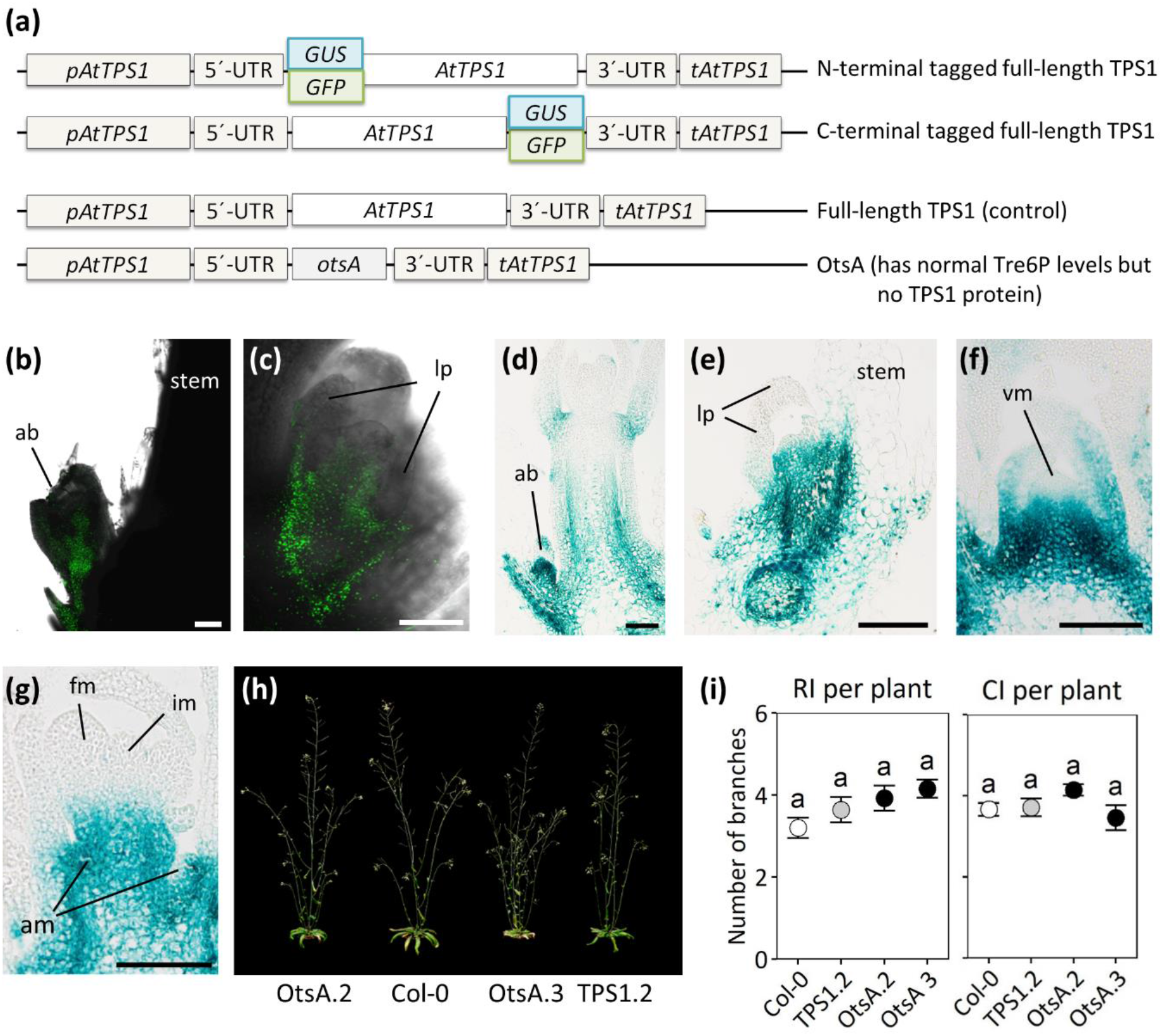
TPS1 localization and Tre6P synthesis during bud development. (a) *TPS1* constructs are derived from the arabidopsis *TPS1* (At1g78580) genomic locus, including the native promoter (*pTPS1*) and terminator (*tTPS1*) regions (Fichtner et al., 2020). TPS1 fusion proteins tagged with the GREEN FLUORESCENT PROTEIN (GFP; green box) or the β-GLUCURONIDASE (GUS; blue box) were used to analyze the TPS1 expression pattern. A construct encoding the full-length TPS1 protein was used as a control together with a construct expressing only the heterologous TPS from *Escherichia coli* (OtsA) under the control of the *TPS1* gene regulatory elements (*pTPS1* and *tTPS1*). Plants were grown in 16-h photoperiods and stained for GUS activity or examined for GFP. (b-c) TPS1-GFP fusion protein expression in axillary buds. (d-e) TPS1-GUS fusion protein expression in rosette axillary buds. TPS1-GUS fusion protein expression in (f) vegetative and (g) inflorescence and floral meristems. (h) Visual phenotype of wild-type Col-0 plants and full-length TPS1 as well as OtsA complemented *tps1-1* plants photographed at 44 DAS. (i) Primary rosette branches (RI) or cauline branches (CI) per plant (length ≥0.5 cm) were counted at the end of the plant’s life cycle. Data are presented as mean ± S.E.M. (*n* = 13-15). Wild-type and transgenic lines complementing the *tps1-1* mutant are represented by different symbol colours: Col-0 (white), full length TPS1 (grey), OtsA (black). Letters indicate significant differences between treatments according to one-way ANOVA with post hoc LSD testing (*p* ≤ 0.05). vm, vegetative meristem; im, inflorescence meristem; fm, floral meristem; am, axillary meristem; ab, axillary bud; lp, leaf primordia; scale bar = 100μm.

The synthesis of Tre6P is the primary function of TPS1. However, some trehalose pathway enzymes are known to have non-catalytic functions as well, including at least two TPP isoforms that influence inflorescence branching in maize (Claeys et al., 2019). To investigate the dependence of shoot branching in arabidopsis on TPS1, we analyzed shoot branching patterns in two independent transgenic lines in the *tps1-1* null mutant background, in which loss of TPS1 had been complemented by expression of the *E. coli* TPS (OtsA) under the control of the *TPS1* promoter and other *TPS1* gene regulatory elements (Fig. 1a). These lines have no detectable TPS1 protein but wild-type levels of Tre6P (Fichtner et al., 2020). Thus, by comparing these with wild-type plants and a *tps1-1* mutant line complemented with *TPS1*, we can determine the dependence of any phenotypes on Tre6P. Both OtsA-complemented lines had the same number of primary rosette (RI) and cauline (CI) branches as wild-type plants and *TPS1*-complemented control plants, showing that Tre6P, rather than the TPS1 protein, is a key factor in regulation of shoot branching (Fig. 1h,i).

Together, these results indicate that there is the enzymatic capacity for Tre6P synthesis in axillary meristems and buds, and that replacement of the Tre6P-synthesizing function of TPS1 by OtsA is sufficient to restore wild-type patterns of shoot branching in the *tps1-1* mutant background.

### Expression of a heterologous TPP in axillary buds supresses branching

To determine whether Tre6P is required for release of axillary bud dormancy and outgrowth into new shoots, we expressed a heterologous TPP in axillary buds to lower their Tre6P content. We used the promoter of the arabidopsis *BRC1* gene, which is predominantly expressed in axillary buds (Aguilar-Martinez et al., 2007), to drive expression of a heterologous TPP from *Caenorhabditis elegans* (CeTPP), whose *K_m_* for Tre6P (0.1-0.15 mM; Kormish and McGhee, 2005) is in a similar range to estimates of *in-vivo* Tre6P concentrations in plants (Martins et al., 2013). CeTPP is phylogenetically unrelated to plant TPPs, so is likely to have unregulated activity when expressed in plants and unlikely to have any physiological interactions with plant proteins. Therefore, any phenotypic effects arising from its expression in axillary buds could be ascribed solely to changes in Tre6P levels. We also generated several independent GUS and GFP reporter lines to confirm the expression pattern mediated by the *BRC1* promoter.

In the inflorescence apex, *pBRC1*-driven GUS expression was visible in axillary meristems and in the epidermis of young leaves (Fig. 2a shows representative images from several independent lines), consistent with the predominant expression in axillary buds described previously (Aguilar-Martinez et al., 2007). Upon bolting, GUS expression was visible in axillary meristems of cauline leaves as well as in axillary buds of rosette leaves (Fig. 2a). Consistent with the GUS expression results, GFP fluorescence was detected in axillary buds of *pBRC1:GFP* lines (Fig. S1a). To confirm transgene expression in the *pBRC1* lines, mature axillary buds and fully expanded leaves were harvested from wild-type Col-0, *pBRC1:GUS* and *pBRC1:CeTPP* lines at one week after bolting for immunoblot analysis (Fig. 2b). In both of the *pBRC1:CeTPP* lines, an immunoreactive protein of the expected size of CeTPP (51 kDa) was detected only in the axillary bud material, with no detectable CeTPP protein in fully expanded leaf tissue (Fig. 2b). This confirms specific expression in mature axillary buds in these lines.

**Fig 2.**
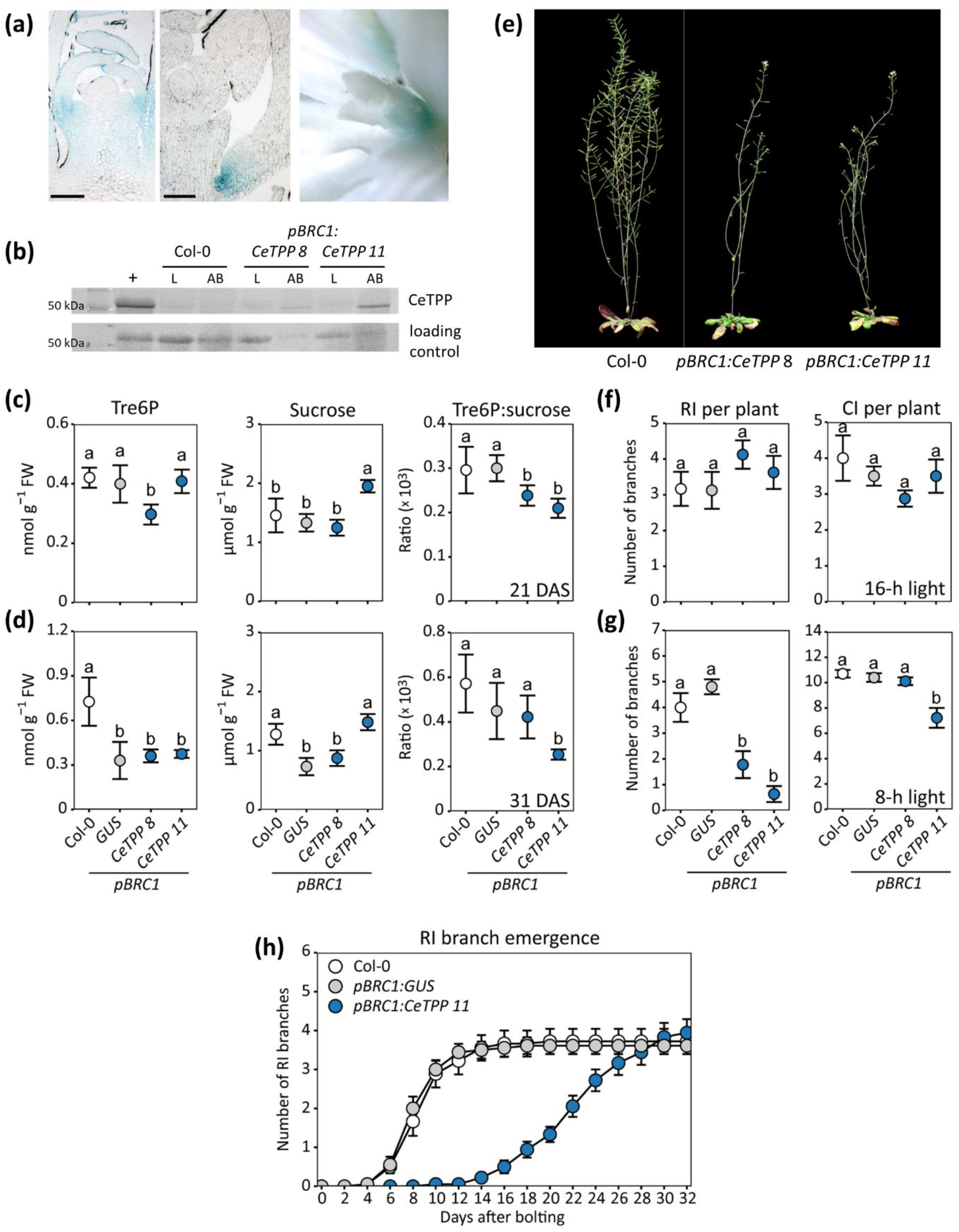
Over-expression of TPP in axillary buds of arabidopsis. The arabidopsis *BRANCHED1* promoter (*pBRC1*) was used to drive specific expression in axillary buds. (a) The β-GLUCURONIDASE (*GUS*) reporter gene was expressed in arabidopsis Col-0 plants under the control of the *pBRC1* promoter and GUS activity was visualized in inflorescence meristems and dormant axillary buds. Scale bar = 200 μM. (b) Wild-type Col-0 and transgenic *pBRC1:GUS,* or *pBRC1:CeTPP* (*Caenorhabditis elegans GOB1*) lines were grown in a 16-h photoperiod. Leaves (L) and axillary buds (AB) were harvested around ZT10 for immunoblotting to detect expression of heterologous CeTPP proteins. Parallel samples of rosette cores were harvested at ZT10 from plants at (c) 21 and (d) 31 DAS for metabolite analyses. Data are presented as mean ± S.D. (*n* = 4). (e) Visual phenotypes of *pBRC1:CeTPP* arabidopsis plants photographed approx. 18 days after bolting. Primary rosette (RI) or cauline (CI) branches were counted (f) in 16-h or (g) 8-h photoperiods. (h) The number of RI branches (length ≥0.5 cm) per plant was counted at 2-d intervals after bolting in a 16-h photoperiod. Branches (length ≥0.5 cm) were counted at the end of the plant’s life cycle. Data are shown as mean ± S.E.M. (*n* = 8-18). Wild-type and transgenic lines expressing heterologous proteins are represented by different symbol colours: Col-0 (white), GUS (grey), and CeTPP (blue). Letters indicate significant differences between treatments according to one-way ANOVA with post hoc LSD testing (*p* ≤ 0.05). +, positive control.

We also measured Tre6P and sucrose levels in tissue samples from rosette cores, enriched in axillary meristems and buds, at various stages of development: at 21 days after sowing (DAS), when the plants had undergone the floral transition and produced axillary meristems, and at 31 DAS, when axillary buds had formed. The level of Tre6P was significantly decreased in *pBRC1:CeTPP* line #8 at 21 DAS, while sucrose was significantly increased in *pBRC1:CeTPP* line #11 at 21 DAS (Fig. 2c). The Tre6P:sucrose ratio was significantly decreased in *pBRC1:CeTPP* line #8 at 21 DAS and in *pBRC1:CeTPP* line #11 at both time points (Fig. 2c,d).

The *pBRC1:CeTPP* plants with lower levels of Tre6P in axillary buds did not show any obvious changes in their vegetative growth pattern (Fig. S2). Similarly, flowering time and final RI branch number were unchanged (Fig. 2e,f; Fig. S4a). However, when branching was scored in short-day conditions (8-h photoperiod), RI number was drastically reduced in the *pBRC1:CeTPP* lines compared to the controls (Fig. 2g). We also analysed the difference in bud outgrowth by monitoring RI branch emergence over time in a 16-h photoperiod. Since independent lines showed consistent phenotypes, we focussed on a single line that showed the strongest accumulation of CeTPP protein (*pBRC1:CeTPP* line #11). The emergence of RI branches was strongly delayed in the *pBRC1:CeTPP* plants, with the first appearance of RI branches occurring about 8 d later than in the controls. The emergence of further RI branches was also noticeably slower in the *pBRC1:CeTPP* plants. As seen before, the final number of RI branches was the same in *pBRC1:CeTPP* and control plants (Fig. 2f,h), indicating that lowering bud Tre6P levels delays the initiation and rate of bud outgrowth, but does not affect the number of buds that eventually do form RI branches in long days. However, lowering Tre6P in axillary buds under carbon limiting conditions (e.g. short days) can successfully inhibit branching in arabidopsis.

### Tre6P levels in the vasculature affect shoot branching in arabidopsis

Arabidopsis is an apoplastic phloem loading species, in which sucrose is released from phloem parenchyma cells into the apoplast via SUCROSE WILL EVENTUALLY BE EXPORTED type sucrose effluxers (SWEET11 and SWEET12) and then actively taken up into the companion cell-sieve element complex by the SUT1/SUC2 sucrose-H^+^ symporter (Lalonde et al., 2003; Baker et al., 2012; Chen et al., 2012; Eom et al., 2015; Zakhartsev et al., 2016). In arabidopsis leaves, the leaf vasculature is a major site of TPS1 expression, and by implication Tre6P synthesis and signalling (Fig. 1; Fichtner et al., 2020). The phloem-loading zone where TPS1 is expressed lies at the interface between source and sink tissues and is therefore a strategically important site for systemic signalling. To investigate the potential of Tre6P synthesis in the vasculature to influence shoot branching, we modified Tre6P levels preferentially in vascular tissues. We used the promoter of the *GLYCINE-DECARBOXYLASE P-SUBUNIT A* (*GLDPA*) gene from the C4 plant *Flaveria trinervia* to drive specific expression of *otsA* or *CeTPP* to increase or decrease Tre6P, respectively, in vascular tissues of arabidopsis. The *GLDPA* promoter had previously been reported to drive specific expression in vascular tissues in arabidopsis (Engelmann et al., 2008; Wiludda et al., 2012; Aubry et al., 2014), which for brevity we shall refer to collectively as the vasculature. We also generated *pGLDPA:GUS* and *pGLDPA:GFP* reporter lines to verify the expression pattern of the *GLDPA* promoter in arabidopsis. For comparison, we expressed *otsA* under the control of the constitutive Cauliflower Mosaic Virus *35S* promoter, as this had previously been reported to give a bushy phenotype (Schluepmann et al., 2003), although no quantitative analysis of shoot branching was reported.

GUS activity was localized to the vascular tissue in the leaves and petioles of *pGLDPA:GUS* lines (Fig. 3a, Fig. S1b), with transverse sections showing staining throughout the vascular bundles (xylem and phloem tissues; Fig. 3a shows representative images from several independent lines). There was expression in the vasculature subtending dormant axillary buds, with a sharp boundary at the base of the bud and no detectable expression in the bud itself (Fig. 3a). Expression was also seen in the vasculature subtending vegetative and inflorescence SAMs but not in the SAM itself (Fig. S1c). In *pGLDPA:GFP* lines, no GFP signal was detected outside of the vasculature (Fig. S1d). To confirm correct transgene expression, immunoblot analyses were performed in extracts of: (i) whole leaves and (ii) dissected mid-veins, the latter being substantially enriched in vascular tissue. The abundance of the OtsA protein in vasculature-enriched extracts from the *p35S:otsA* plants was similar to that in whole-leaf extracts (Fig. 3b), indicating expression at a similar level throughout the leaf. In contrast, OtsA and CeTPP were more abundant in the vasculature-enriched samples from two independent *pGLDPA:otsA* and *pGLDPA:CeTPP* lines, respectively, than in the corresponding whole-leaf extracts (Fig. 3b). Together, the GUS/GFP reporter lines and the immunoblotting data confirm that the *GLDPA* promoter drives vasculature specific expression of heterologous proteins in arabidopsis.

**Fig 3.**
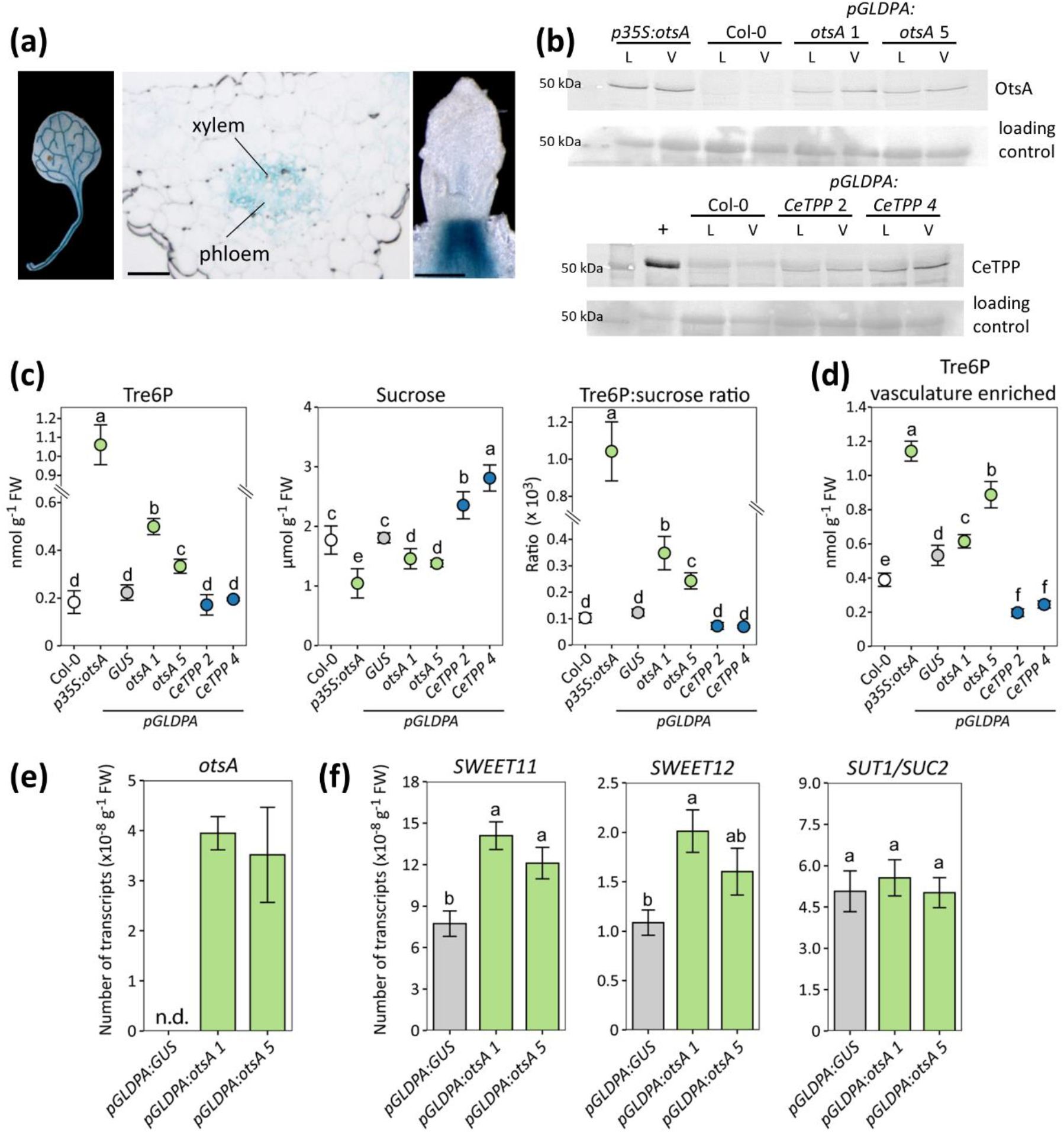
Over-expression of TPS or TPP in the vasculature of arabidopsis. The *Flaveria trinervia GLYCINE-DECARBOXYLASE P-SUBUNIT A* promoter (*pGLDPA*) was used to drive specific expression in the vasculature of Arabidopsis. (a) The *β-GLUCURONIDASE* (*GUS*) reporter was expressed in arabidopsis Col-0 plants under the control of *pGLDPA* and GUS activity was visualized in rosette leaves, transverse sections of the mid-vein region of fully expanded leaves and in dormant axillary buds. Scale bar = 100μm. (b) Wild-type Col-0 and transgenic *p35S:otsA* (*Escherichia coli otsA*), *pGLDPA:GUS*, *pGLDPA:otsA* and *pGLDPA:CeTPP* (*Caenorhabditis elegans GOB1*) lines were grown in a 16-h photoperiod. Leaves (L) and vasculature-enriched (V) tissues were harvested at ZT10 from plants at 20 DAS to detect expression of heterologous TPS (OtsA) or TPP (CeTPP) proteins by immunoblotting. Parallel samples of whole rosettes (c) or vasculature-enriched tissue (d) were collected for metabolite analyses. Data are presented as mean ± S.D. (*n* = 4). (e) Transcript abundance of the *otsA* gene in rosettes from plants grown under long-day conditions and harvested at ZT10 at 24 DAS. (F) Transcript abundance of *SUT1*/*SUC2, SWEET11* and *SWEET12* in the same samples as (E). Data are presented as mean ± S.E.M. (*n* = 4 biological replicates). Wild-type and transgenic lines expressing heterologous proteins are represented by different symbol colours: Col-0 (white), GUS (grey), OtsA (green) and CeTPP (blue). Letters indicate significant differences between treatments according to one-way ANOVA with post hoc LSD testing (*p* ≤0.05). +, positive control; n.d., not detected.

Tre6P was significantly increased in rosettes of the *p35S:otsA* plants (5-fold) as well as in both *pGLDPA:otsA* lines (2-fold) compared to wild-type Col-0 and control plants (Fig. 3c). The *p35S:otsA* and *pGLDPA:otsA* lines had lower sucrose levels, resulting in a significantly increased Tre6P:sucrose ratio in the *p35S:otsA* (10-fold) plants and in both *pGLDPA:otsA* lines (3-fold; Fig. 3c). The *pGLDPA:CeTPP* lines had significantly higher rosette sucrose levels (Fig. 3c). In vasculature-enriched samples, Tre6P levels were significantly higher than wild-type in *p35S:otsA* and *pGLDPA:otsA* lines, while the *pGLDPA:CeTPP* lines had significantly less Tre6P (30-40% lower than wild-type; Fig. 3d).

Tre6P has been implicated in transcriptional regulation of *SWEET* expression in sorghum and maize (Kebrom and Mullett, 2016; Bledsoe et al., 2017; Oszvald et al., 2018). Therefore, we measured the transcript abundance of *SWEET* and other sucrose transporter (*SUT*/*SUC*) genes to investigate the potential impact of increased Tre6P levels in the vasculature on phloem loading of sucrose. Expression of *otsA* was confirmed in both *pGLDPA:otsA* lines, with no *otsA* transcript being detected in the *pGLDPA:GUS* control line (Fig. 3e). *SUT1*/*SUC2* transcript abundance was the same in all genotypes, but the two *pGLDPA:otsA* lines had 1.5-1.8 times higher levels of *SWEET11* and *SWEET12* transcripts (Fig. 3f).

To follow up these preliminary observations, we measured the transcript abundance of all four *SWEET* genes (*SWEET11*-*SWEET14*) that are known to encode plasmalemma sucrose efflux carriers (Chen et al., 2012; Kanno et al., 2016) in leaf 6 (a fully expanded source leaf) and in the rosette core (enriched in axillary buds) from *pGLDPA:otsA* (line #5) and control *pGLDPA:GUS* plants at 2 d after bolting (Fig. 4a). *SWEET11* and *SWEET12* were up-regulated about 2.5-fold in the *pGLDPA:otsA* plants (Fig. 4a), consistent with our previous results based on whole rosette measurements that were performed at a different biological age and time of the day (Fig. 3f). *SWEET13* was also significantly up-regulated (3-fold), but *SWEET14* transcripts were not detected (Fig. 4a). In contrast to leaves, there was no up-regulation of *SWEET11* and *SWEET12* in axillary bud enriched rosette cores but transcripts of *SWEET13* and *SWEET14* were 3-times more abundant in rosette cores from the *pGLDPA:otsA* plants than in the controls (Fig. 4a).

**Fig 4.**
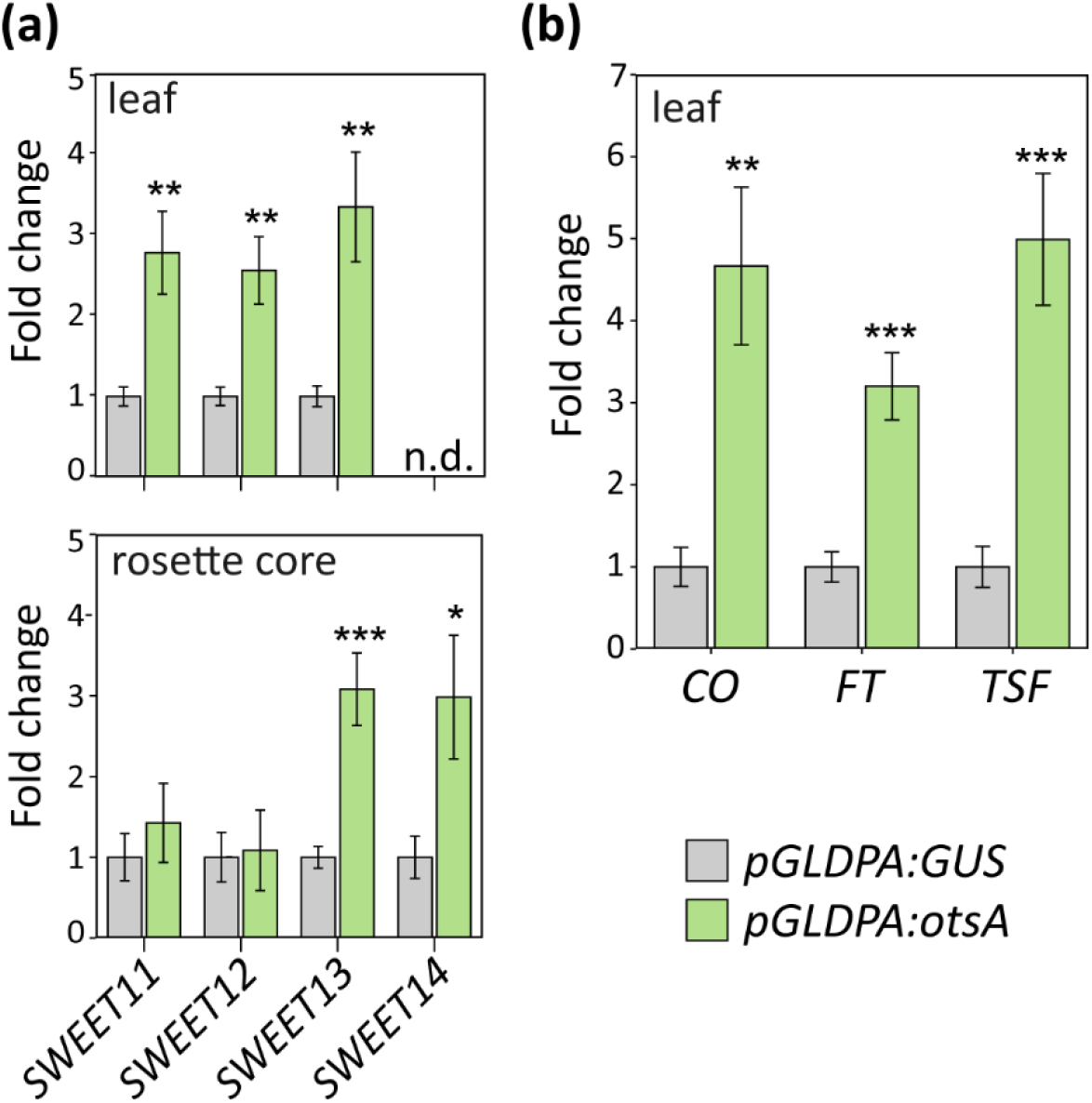
Gene expression analyses in arabidopsis plants with altered Tre6P levels in the vasculature. *pGLDPA:otsA* (green bars) and *pGLDPA:GUS* (grey bars) plants were grown in a 16-h photoperiod. Fully expanded leaves (leaf 6) and rosette cores were harvested at ZT 14 on day 2 after bolting, i.e. after the floral transition but before axillary bud outgrowth. (a) Abundance of *SWEET11*, *SWEET12*, *SWEET13* and *SWEET14* transcripts. (b) Abundance of *CONSTANS* (*CO*), *FLOWERING LOCUS T* (*FT*) and *TWIN SISTER OF FLOWERING LOCUS T* (*TSF*). Data from the *pGLDPA:otsA* plants are expressed as fold-change with respect to the *pGLDPA:GUS* controls, and shown as mean ± S.E.M. (*n* = 8 biological replicates). Asterisks show significant differences between the genotypes according to Student’s t-test: * *p* < 0.05, ** *p* < 0.01, *** *p* < 0.001. n.d., not detected.

As previously observed (Schluepmann et al., 2003; Yadav et al., 2014), the *p35S:otsA* plants had much smaller rosettes and darker green leaves than wild-type Col-0 plants (Fig. S3). Similarly, the *pGLDPA:otsA* lines had smaller rosettes than wild-type, although not as small as the *p35S:otsA* plants, whereas the *pGLDPA:CeTPP* lines had slightly bigger leaves than wild-type (Fig. S3).

In long-day conditions *p35S:otsA* and *pGLDPA:otsA* plants displayed increased shoot branching (Fig. 5a). On average, wild-type Col-0 and control plants developed only two to three RI branches, while the *p35S:otsA* and *pGLDPA:otsA* lines had four to six RI branches (Fig. 5b). RI branch number in the *pGLDPA:CeTPP* lines was similar to wild-type Col-0 and control plants (Fig. 5b). In arabidopsis, an axillary bud develops in the axil of each rosette leaf. Therefore, the number of RI branches per plant is potentially influenced by the number of rosette leaves at the floral transition, when the plant stops initiating new rosette leaves. Leaf number also influences the overall photosynthetic capacity and potential sucrose supply of the plant. As total leaf number was decreased in the *p35S:otsA* and *pGLDPA:otsA* lines and increased in the *pGLDPA:CeTPP* lines (Fig. S4c), the total number of RI branches was also plotted on a per rosette leaf basis (RI per leaf). The *p35S:otsA* plants and both *pGLDPA:otsA* lines had twice as many RI per leaf as wild-type and control plants (Fig. 5b), whereas *pGLDPA:CeTPP* line #4 showed the opposite phenotype, with only half the wild-type number of RI per leaf (Fig. 5b). The stronger phenotype of *pGLDPA:CeTPP* line #4 compared to line #2 was consistent with the higher abundance of CeTPP protein in this line (Fig. 3b). In addition to the increase in RI, the *pGLDPA:otsA* lines also had a significantly increased ratio of secondary (RII) to primary rosette branches (RII:RI ratio; Fig. 5b), indicating that outgrowth of secondary buds was also stimulated in these plants. We also determined the number of CI per plant. As the *pGLDPA:otsA* lines flowered earlier than the controls, they produced fewer CI, whereas the late flowering *pGLDPA:CeTPP* plants developed more CI (Fig. 5b).

**Fig 5.**
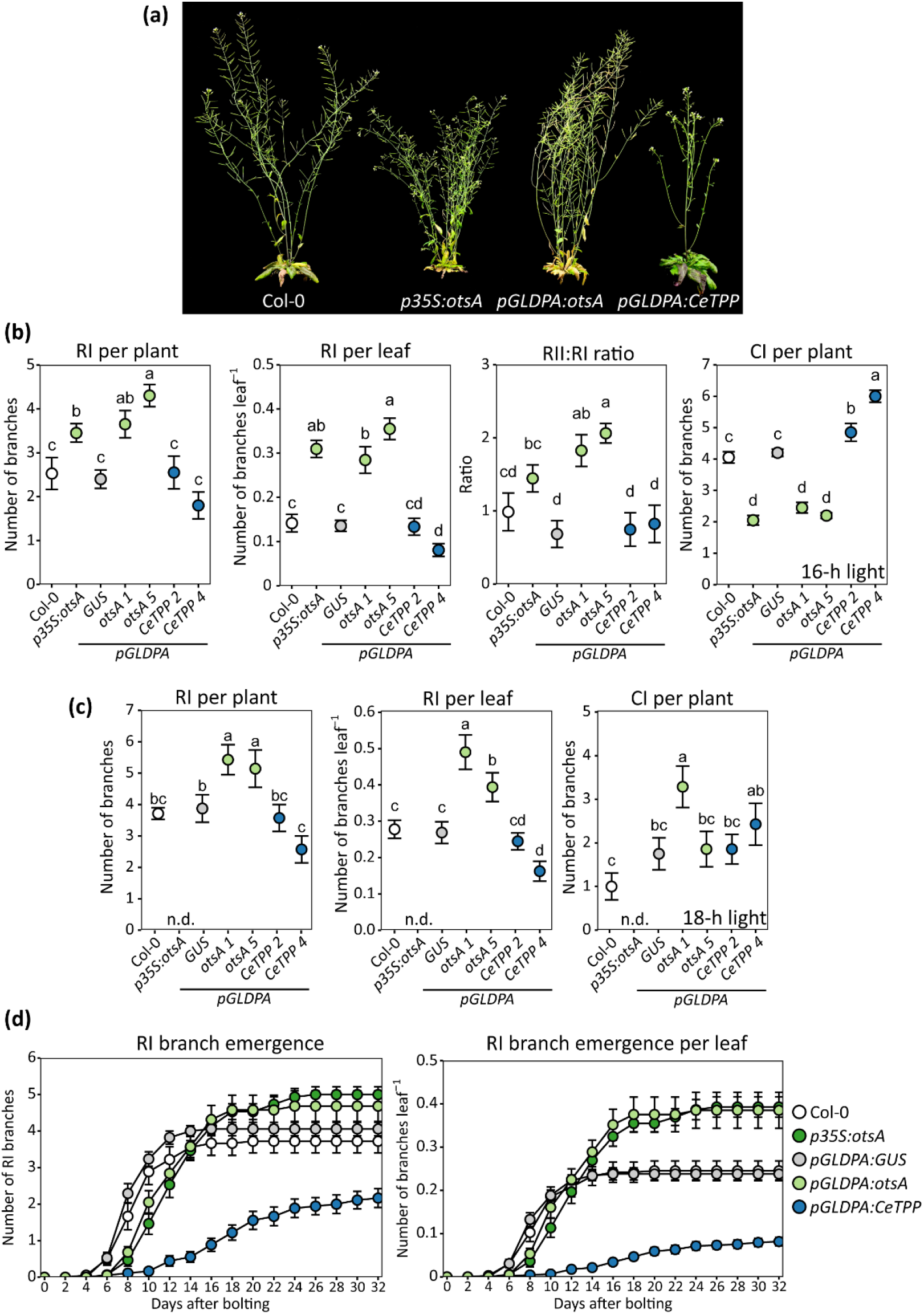
Axillary bud growth analysis in arabidopsis plants with altered Tre6P levels in the vasculature. (a)Visual phenotypes of arabidopsis *p35S:otsA, pGLDPA:otsA* and *pGLDPA:CeTPP* plants, grown in a 16-photoperiod and photographed at 56 DAS. Primary rosette branches (RI) per plant, RI branches per rosette leaf, ratio of secondary (RII) and RI branches and primary cauline branches (CI) per plant were counted in (b) 16-h or (c) 18-h photoperiods. (d) The number of RI branches per plant and per leaf was counted at 2-d intervals after bolting of plants grown in a 16-h photoperiod (n.b. the wild-type data are from plants grown under identical conditions and are the same as those shown in Fig.2h). Branches (length ≥0.5 cm) were counted at the end of the plant’s life cycle. Data are shown as mean ± S.E.M. (*n* = 8-20). Wild-type and transgenic lines expressing heterologous proteins are represented by different symbol colours: Col-0 (white), GUS (grey), OtsA (green) and CeTPP (blue). Letters indicate significant differences between genotypes according to one-way ANOVA with post hoc LSD testing (*p*≤0.05). n.d., not determined.

Essentially the same effects on shoot branching and leaf number were observed in an independent experiment where the plants were grown in a slightly longer (18 h) photoperiod (Fig. 5c, S4d). However, when grown in an 18-h photoperiod, total leaf number was similar for *pGLDPA:otsA* and control plants (Fig. S4d), and there were no differences in CI number, except for *pGLDPA:otsA* line #1 which had more CI due to formation of a second CI branch at some axils (Fig. S5).

To determine if there is also a difference in the rate of branch emergence, the number of RI was monitored over time in long-day grown plants and plotted against days after bolting. Initiation of new branches continued for longer in the *p35S:otsA* and *pGLDPA:otsA* (line #5) plants, so by the time the plants were fully senesced they had more RI per plant and RI per rosette leaf than the control plants (Fig. 5d). In the *pGLDPA:CeTPP* (line #4) plants, the emergence of lateral branches was slower than in the control plants (Fig. 5d).

As both flowering time and branching were affected in the *pGLDPA:otsA* plants with elevated Tre6P in the vasculature, we compared the expression levels of *FT* and its close homologue, *TSF*, in *pGLDPA:otsA* plants with those in a *pGLDPA:GUS* control line. We also included *CONSTANS* (*CO*) in the analysis as this is a key regulator of *FT* expression (Turck et al., 2008). Transcript analysis was performed on the same RNA samples from leaf 6 harvested from *pGLDPA:otsA* and *pGLDPA:GUS* plants grown in a 16-h photoperiod as described above (Fig. 4b). Compared to the *pGLDPA:GUS* controls, the *pGLDPA:otsA* plants showed significantly increased expression of *CO* (4.5-fold), *FT* (3-fold) and *TSF* (5-fold) (Fig. 4b).

To summarize, the phenotypes of these transgenic lines demonstrate that: (i) higher Tre6P in the vasculature (*pGLDPA:otsA*) increased the final number of branches and was associated with up-regulation of *SWEETs, CO*, *FT* and *TSF*; and (ii) lowering Tre6P in the vasculature (*pGLDPA:CeTPP*) decreased final branch number and strongly delays lateral bud outgrowth.

### Rosette branch number correlates with Tre6P levels in the vasculature

To confirm that the level of Tre6P correlates with bud outgrowth, we assessed the relationship between Tre6P, sucrose and the Tre6P:sucrose ratio in whole-rosette or vascular-enriched samples with the number of RI branches per plant and RI branches per leaf, using a Pearson’s correlation analysis. Based on whole rosette measurements, we detected a significant negative correlation of sucrose levels with RI per plant and with RI per leaf (RI per plant: r = −0.813, R^2^ = 0.66, *p*<0.026; RI per leaf: r = −0.855, R^2^ = 0.732, *p*<0.014). The manipulation of Tre6P in the vasculature influences the rosette’s sucrose levels (see also Fig. 3c) resulting in the negative correlation between sucrose levels and rosette branching. There was no significant correlation between Tre6P or the Tre6P:sucrose ratio based on whole rosette measurements. However, based on measurements of vascular enriched samples, both Tre6P and the Tre6P:sucrose ratio had a highly significant positive correlation with RI number per plant and an even stronger positive correlation with RI number per leaf (Fig. 6). Thus, theTre6P:sucrose ratio in the vasculature correlates best with the observed branching phenotype in the *pGLDPA* lines.

**Fig 6.**
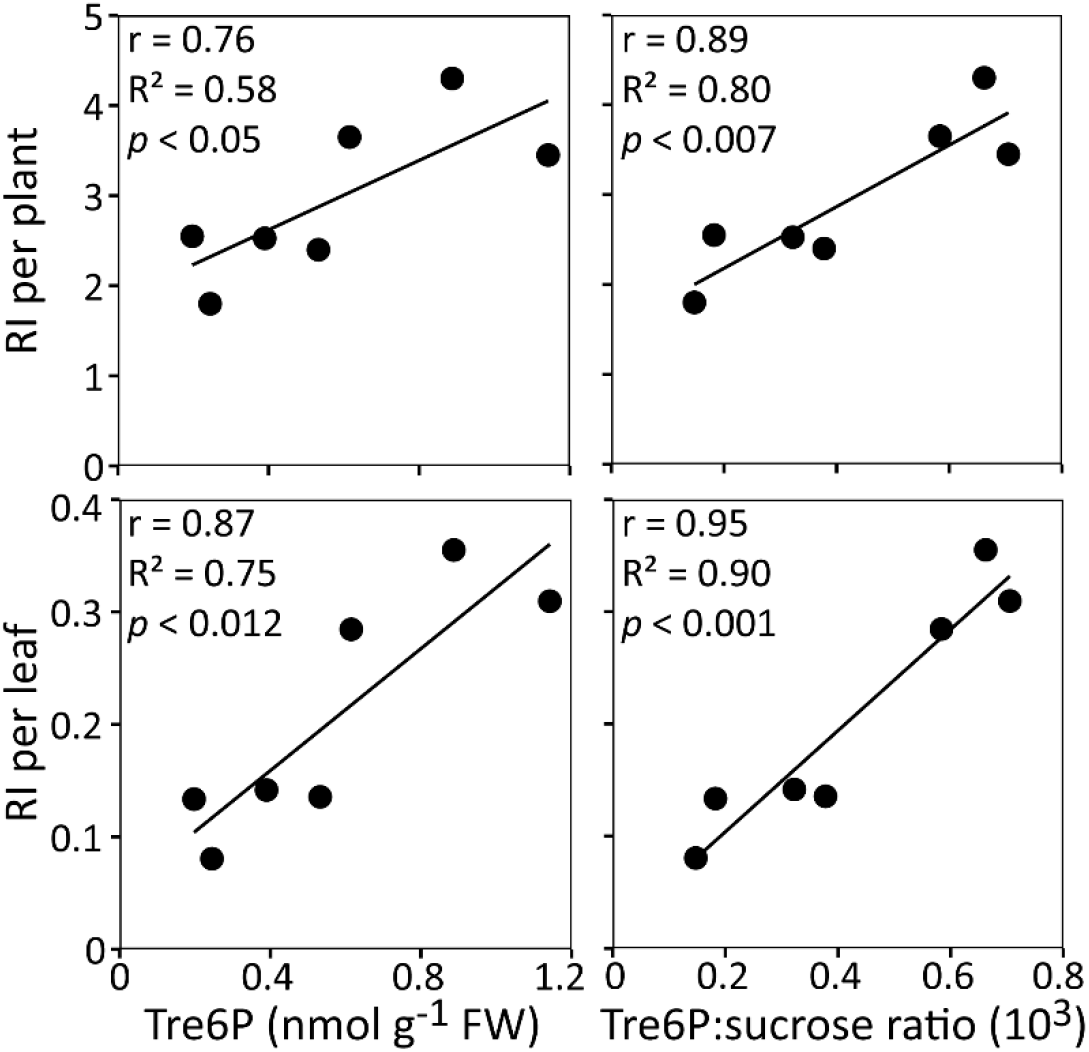
Correlation analyses in arabidopsis plants with altered Tre6P levels in the vasculature. (A)The number of RI branches per plant and per leaf of *pGLDPA:otsA*, *pGLDPA:CeTPP*, *pGLDPA:GUS*, *p35S:otsA* and wild-type Col-0 plants were plotted against the level of Tre6P and the Tre6P:sucrose ratio measured in vasculature-enriched tissue samples. Plants were grown in 16-h photoperiods. The Pearson correlation coefficient (r), coefficient of determination (R^2^) and probability (*p*) values for each relationship are indicated.

### Interaction between Tre6P and the BRC1 branching signal integrator

To investigate how Tre6P signalling in the vasculature might be integrated with other signalling pathways that regulate shoot branching, *pGLDPA:otsA* (line #1) was crossed with the *brc1-2* (*brc1*) mutant. Given the potential role of FT in branching, the up-regulation of *FT* in the *pGLDPA:otsA* plants and their branching phenotype, we also crossed *pGLDPA:otsA* (line #1) with the *flowering locus t-10* (*ft*) mutant and analysed RI branch number of plants grown in a 16-h photoperiod.

Compared with wild-type Col-0 plants, *pGLDPA:otsA* plants had more branches (on average 3.6 RI branches) than the wild-type (on average 2.5 RI branches; Fig. 7a,c), confirming our previous results. The total number of RI was also increased in *brc1* (on average nine RI branches) and increased still further in *brc1* x *pGLDPA:otsA* double mutant plants (on average 15 RI branches; Fig. 7a,c). In contrast, branching was almost abolished in both the *ft* and *ft* x *pGLDPA:otsA* mutants (on average fewer than one RI branch; Fig. 7b,c). As seen before, *pGLDPA:otsA* plants flowered earlier than wild-type plants, as did the *brc1* x *pGLDPA:otsA* and *ft* x *pGLDPA:otsA* lines compared to their respective *brc1* and *ft* parents (Fig. S6). Given the differences in total leaf number, we also plotted RI on a per rosette leaf basis. The *brc1* x *pGLDPA:otsA* double mutant had the highest number of RI branches/leaf (1.0; Fig. 7c), indicating that every axillary rosette bud grew out in this line. The ratio of RI per rosette leaf was lower in the *brc1* mutant (0.5), the *pGLDPA:otsA* line (0.3) and wild-type plants (0.15). The *ft* mutant (0.01) and the *ft-10* x *pGLDPA:otsA* double mutant (0.02) had the lowest values. We confirmed these results in a second independent experiment with the same lines (Fig. S7).

**Fig 7.**
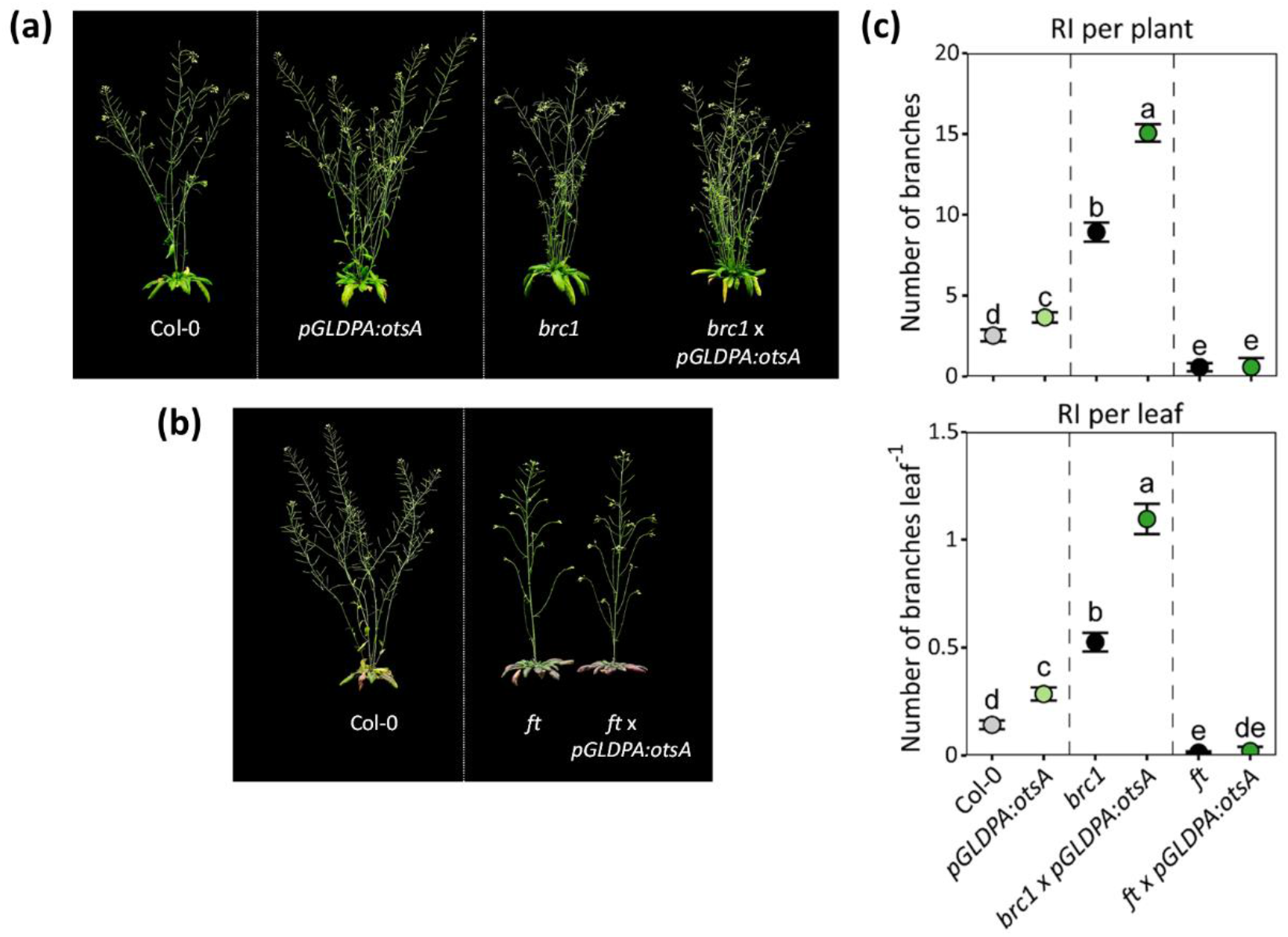
Effects of vasculature-specific TPS overexpression in wild-type and branching mutant backgrounds on flowering and shoot branching under long-day conditions. Visual phenotypes of (a) *branched1* (*brc1*) and (b) *flowering locus t* (*ft*) mutants and plants expressing the *Escherichia coli* TPS (OtsA) in the vasculature (*pGLDPA:otsA*) in these mutant backgrounds grown in a 16-h photoperiod. Plants were grown in a 16-h photoperiod and photographed at 45 (a) or 56 (b) DAS. (c) Primary rosette branches (RI) per plant, flowering time based on total leaf number and RI branches per rosette leaf. Symbol colours: wild-type Col-0 (grey); *brc1*, *ft* parental mutants (black); *pGLDPA:otsA* expression in a wild-type background (light green); *pGLDPA:otsA* expression in a mutant background (dark green). RI branches (length ≥0.5 cm) were counted at the end of the plant’s life cycle. Data are presented as mean ± S.E.M. (*n* = 11-19) and letters indicate significant differences between genotypes according to one-way ANOVA with post hoc LSD testing (*p* ≤ 0.05).

In summary, the branching mutant analysis showed that loss of BRC1 and increased Tre6P in the vasculature had a strong additive effect on shoot branching, while increasing Tre6P in the vasculature in the *ft* mutant background had no impact on axillary bud outgrowth.

## DISCUSSION

### Tre6P acts locally within axillary buds to modulate shoot branching

We previously demonstrated that the breaking of axillary bud dormancy in pea is associated with a rapid rise in bud Tre6P levels that is highly correlated with bud outgrowth (Fichtner et al. 2017). In arabidopsis, we observed that the predominant Tre6P-synthesizing enzyme, TPS1, is expressed in axillary buds, and that complementation of the *tps1-1* mutant by expression of a heterologous TPS from *E. coli* (OtsA) restored shoot branching to wild-type levels (Fig. 1). Together, these results show that there is enzymatic capacity for Tre6P synthesis in axillary buds and that the influence of TPS1 on shoot branching is primarily due to its Tre6P-synthesizing activity rather than any non-catalytic (e.g. signalling) function. With TPS1 being present in the buds, we can infer that Tre6P is at least partly produced locally within the buds in response to any increase in their sucrose supply. Expressing CeTPP in buds to counteract any rise in their Tre6P levels led to delayed rosette branching in the *pBRC1:CeTPP* plants in long-days (Fig. 2h) and the suppression of branching in short days (Fig. 2g), providing genetic evidence that Tre6P acts locally within axillary buds to modulate shoot branching.

### Tre6P levels in the vasculature regulate shoot branching in arabidopsis

The expression pattern of the *F. trinervia GLDPA* promoter is very similar to the expression domain of the *TPS1* gene in the vasculature (Fichtner et al., 2020), providing a means to investigate the specific functions of Tre6P in the vasculature. The *pGLDPA:otsA* lines, with increased Tre6P levels only in the vasculature, displayed an increased branching phenotype to the same degree as *p35S:otsA* plants. Our observation that the *pGLDPA:CeTPP* plants had the opposite phenotype provides compelling evidence that these phenotypic differences were driven by changes in Tre6P. Furthermore, there was a strong correlation between branching and the levels of Tre6P and Tre6P:sucrose ratio in the vasculature. Given that TPS1 is expressed predominantly in the vasculature (Fichtner et al., 2020) and the *pGLDPA:otsA* plants flowered early and had as many branches as *p35S:otsA* plants, we conclude that the vasculature is a primary location for Tre6P synthesis and signalling in the regulation of flowering and branching.

### Potential mechanisms for regulation of shoot branching by Tre6P in the vasculature and in axillary buds

Altering Tre6P levels in the vasculature could affect shoot branching in several ways. Tre6P produced in the companion cell-sieve element complex of the vasculature is likely to move with the mass flow of solutes in the phloem and be delivered to distal sink organs, such as axillary buds, where it might supplement Tre6P made locally by the TPS1 enzyme in the buds (Fig. 8). In principle, grafting experiments with *tps1* null mutants would be the simplest approach for testing whether Tre6P does move from source to sink organs via the phloem, but such experiments are not feasible due to the embryo lethality of *tps1* null mutants (Eastmond et al., 2002).

**Fig 8.**
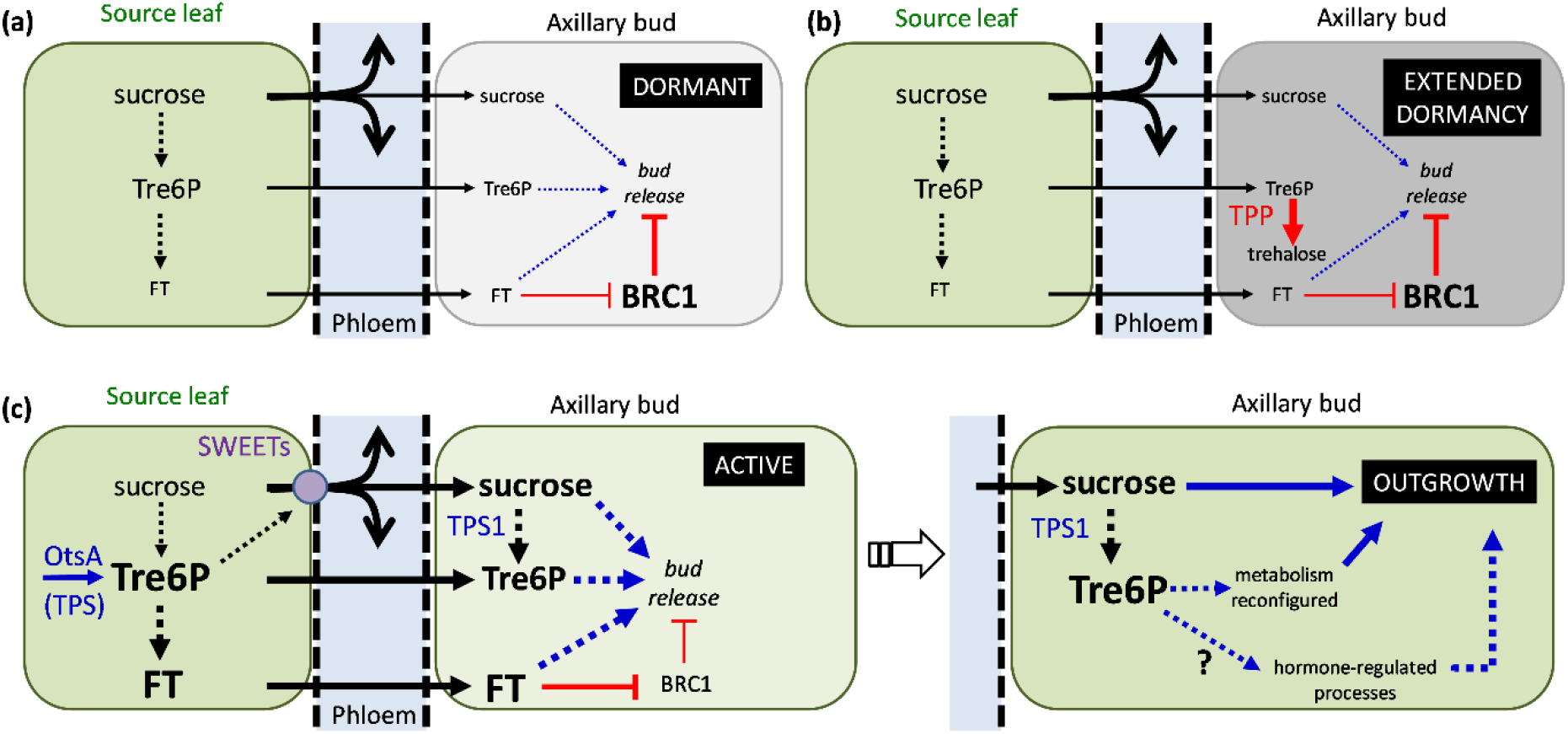
Schematic model for the role of Tre6P in axillary bud outgrowth in arabidopsis. (a) Limited supply of sucrose to axillary buds and strong expression of BRANCHED1 (BRC1) maintains bud dormancy. (b) Bud-specific expression of a heterologous TPP in *pBRC1:CeTPP* lines lowers Tre6P levels in the buds, further delaying their release from dormancy. (c) Expression of a heterologous TPS (OtsA) in *pGLDPA:otsA* lines increases Tre6P in the phloem parenchyma and companion cell-sieve element complex in leaf veins, leading to up-regulation of SWEET sucrose efflux carries, enhanced phloem loading of sucrose and increased sucrose supply to axillary buds. Higher sucrose stimulates local synthesis of Tre6P in the buds by TPS1 (additional Tre6P might also move from leaves to buds via the phloem). In parallel, high Tre6P in companion cells stimulates expression of *FLOWERING LOCUS T* (*FT*). Movement of the phloem-mobile FT protein to buds leads to inhibition of BRC1. High sucrose, high Tre6P and FT act synergistically to trigger the release of bud dormancy. Following release from dormancy, Tre6P sustains bud outgrowth by coordinating a reconfiguration of bud metabolism for growth and via interaction with hormone-regulated processes (e.g. stimulating establishment of polar auxin transport from the new shoot).

Tre6P could also have indirect effects in the vasculature by influencing sucrose allocation (Fig. 8). For example, the observed up-regulation of *SWEET11*, *SWEET12* and *SWEET13* in source leaves in *pGLDPA:otsA* plants (Fig. 3f,4a) could increase the export of sucrose from source leaves and this the potential supply of sucrose to sink organs, including axillary buds. SWEET-type sucrose efflux carriers are also involved in phloem unloading in many sink organs (Eom et al., 2015; Milne et al., 2017) and *SWEET13* and *SWEET14* are expressed in or nearby axillary buds (Kanno et al., 2016). Therefore, the observed up-regulation of *SWEET13* and *SWEET14* in rosette cores of the *pGLDPA:otsA* plants could indicate increased capacity to deliver sucrose to axillary buds and the SAM. Increasing the supply of sucrose to axillary buds via such mechanisms would not only provide more carbon and energy for growth, but could also trigger their release from dormancy and growth into new shoots through a signalling pathway (Mason et al., 2014). In accordance, lowering Tre6P in developing maize seeds led to increased yield under well-watered or drought conditions that correlated with upregulation of *SWEET* transporter genes (Nuccio et al, 2015; Oszvald et al., 2018). Overexpression of SWEETs also leads to more axillary growth in *Chrysanthemum morifolium* (Liu et al., 2020) and early flowering in arabidopsis (Andrés et al., 2020).

A third possibility is that altered Tre6P levels in the vasculature affect other systemic signalling components and pathways that influence shoot branching. In arabidopsis, the expression of *FT* is triggered by long days, under the control of CO, and is dependent on the Tre6P-synthesizing activity of TPS1 (Wahl et al., 2013; Fichtner et al., 2020). FT can also move via the phloem to axillary meristems, and promote their elongation and development by activating their floral transition (Niwa et al., 2013; Tsuji et al., 2015). Plants with higher Tre6P in the vasculature have early flowering and increased branching phenotypes, and we observed significant increases in *CO*, *FT* and *TSF* transcript abundance in the *pGLDPA:otsA* plants (Fig. 4b). This suggested that increased Tre6P levels in the vasculature induced *CO* expression, thereby increasing expression of *FT* and *TSF*, which in turn could result in early flowering and increased branching (Fig. 8). Accordingly, stimulation of branching by increased Tre6P in the vasculature was abolished in an *ft* mutant background (Fig. 7b,c), suggesting that FT is a crucial factor in the ability of axillary buds to respond to distal changes in Tre6P levels (Fig. 8).

We also showed that increasing Tre6P in a *brc1* mutant background has a strong additive effect on branching. This suggests that the stimulation of FT expression in the leaf vasculature and loss of the FT repressor in axillary buds (i.e. BRC1) act synergistically to bring about the strong branching phenotype of the *brc1* x *pGLDPA:otsA* plants. In arabidopsis and wheat, the FT protein (also TSF in arabidopsis) has been shown to interact directly with BRC1, and this interaction leads to a reciprocal repressive effect, i.e. FT and BRC1 inhibit each other (Niwa et al., 2013; Dixon et al., 2018). Enhanced *FT* expression in the *pGLDPA:otsA* lines could lead to an accumulation of FT protein in the buds shifting the FT:BRC1 ratio in favour of FT leading to the stimulation of bud outgrowth (Fig. 8). It was recently reported that a potato (*Solanum tuberosum*) tuber-specific isoform of FT (StSP6A) can interact with StSWEET11 to block the leakage of sucrose to the apoplast and promote symplastic transport of sucrose (Abelenda et al., 2019). Such switches in the pathway of sucrose delivery are a common feature during the development and growth of sink organs (Weber et al., 1988; Eom et al., 2015; Milne et al., 2017). Thus, distal changes in Tre6P could affect the delivery of sucrose to axillary buds via changes in *SWEET* expression, or as described above, via FT-mediated changes in the pathway of phloem unloading in the buds, or both.

Lowering bud Tre6P levels, by *pBRC1*-driven expression of CeTPP, delayed bud outgrowth in long days and suppressed branching in short days, providing genetic evidence that Tre6P acts locally within axillary buds to modulate shoot branching. This suggests that Tre6P, either by itself or by potentiating the effect of other signals (e.g. phytohormones), is involved in modulating bud outgrowth. Low bud Tre6P levels could also compromise Tre6P-driven changes in central metabolism that are needed for sustained outgrowth (Fichtner et al., 2017). In addition, there are several lines of evidence that Tre6P associated changes in bud metabolism might also have an impact on auxin synthesis and signalling. Tre6P promotes the expression of the auxin biosynthesis gene *TRYPTOPHAN AMINOTRANSFERASE RELATED2* in pea seeds (Meitzel et al., 2019) and PINOID1 (PIN1) auxin efflux proteins become delocalized in meristems when plants have low C status (Lauxmann et al., 2016) and therefore low Tre6P levels (Lunn et al., 2006).

In conclusion, we provide genetic evidence that Tre6P acts locally in axillary buds, showing that Tre6P in buds is not only correlated with growth (Fichtner et al., 2017), but also necessary for bud outgrowth. Our results also implicate Tre6P in systemic regulation of bud outgrowth, acting in the vasculature to signal the overall sucrose status of the plant and control sucrose allocation, and being linked to photoperiod signalling by FT under long-day conditions. We postulate that Tre6P is a key factor linking shoot branching to carbon availability, enabling plants to sense and allocate their carbon resources to an appropriate number of shoot branches, thus helping to optimize shoot architecture for survival and reproductive success. In future experiments, the level of Tre6P in the vasculature and axillary buds could be a target for engineering improvements in crop architecture.

## SUPPORTING INFORMATION

**Fig. S1** Tissue specific expression patterns of the *GLDPA* and the *BRC1* promoters in arabidopsis.

**Fig. S2** Rosette morphology of arabidopsis plants expressing a heterologous TPP under the control of an axillary bud-specific promoter (long-day conditions).

**Fig. S3** Rosette morphology of arabidopsis plants expressing heterologous TPS or TPP under the control of constitutive or vasculature-specific promoters (long-day conditions).

**Fig. S4** Flowering time of TPS and TPP over-expression lines.

**Fig. S5** Cauline branching phenotype of arabidopsis plants expressing a heterologous TPS under the control of constitutive or vasculature-specific promoters.

**Fig. S6** Effects of vasculature-specific TPS overexpression in wild-type and branching mutant backgrounds on flowering time under long-day conditions.

**Fig. S7** Effects of vasculature-specific TPS overexpression in wild-type and branching mutant backgrounds on flowering and shoot branching under long-day conditions (independent experiment).

## ACKNOWLEDGEMENTS

We thank Ursula Krause for her excellent help with plant work and immunoblotting, Prof. Dr Peter Westhoff and his colleagues for providing the *FtGLDPA* promoter, Dr Carlos Figueroa for providing the anti-CeTPP antibody, Prof. Pilar Cubas for providing the *brc1-2* mutant, and Dr Elizabeth Dun for helpful comments on the manuscript. This work was supported by a PhD scholarship from the International Max Planck Research School – Primary Metabolism and Plant Growth (F.F.), the Australian Research Council (F.F.B.; Discovery grant DP150102086 and Georgina Sweet Laureate Fellowship FL180100139 to C.A.B.), the Max Planck Society (F.F., M.G.A., R.F., M.S. and J.E.L.) and Deutsche Forschungsgemeinschaft (DFG; within the Collaborative Research Centre 973; J.J.O.; B.M.-R.).

## AUTHOR CONTRIBUTIONS

C.A.B., M.S. and J.E.L conceived the project. F.F. designed and performed all experiments and measurements with help from F.F.B. and M.G.A. R.F. performed Tre6P measurements. J.J.O. and B.M.-R. performed sectioning and helped with imaging of stained sections. F.F. and J.E.L wrote the manuscript with help from F.F.B., M.S. and C.A.B. All authors commented on the manuscript and approved the final version.

## Notes

### Competing Interest Statement

The authors have declared no competing interest.

