## Supplementary Information for "The role of trehalose 6-phosphate in shoot branching – local and non-local effects on axillary bud outgrowth in arabidopsis rosettes"

(a)

(b)

(c)

(d)

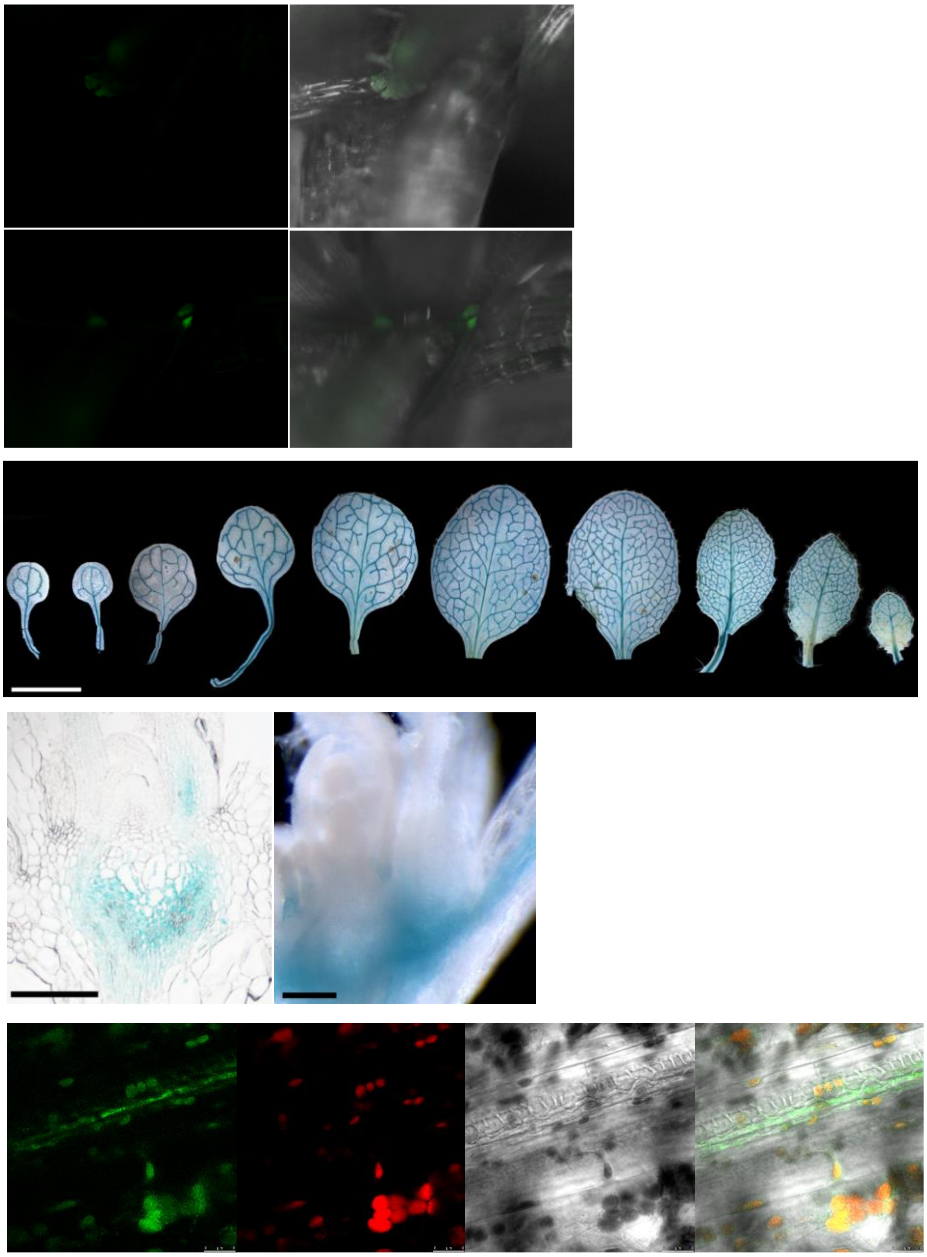

**Fig. S1** Tissue specific expression patterns of the *GLDPA* and the *BRC1* promoters in Arabidopsis. Supports Fig. 1.

(a) Localization of GFP in leaf axils of *pBRC1:GFP* plants. (b) Localization of GUS activity in rosette leaves of *pGLDPA:GUS* plants at 18 DAS (oldest to youngest leaf shown from left to right). Scale bar = 1 mm. (c) Localization of GUS activity in vegetative and inflorescence shoot apical meristems of *pGLDPA:GUS* plants. Scale bar 200 µM. (d) Localization of GFP in leaves of *pGLDPA:GFP* at 12 DAS.

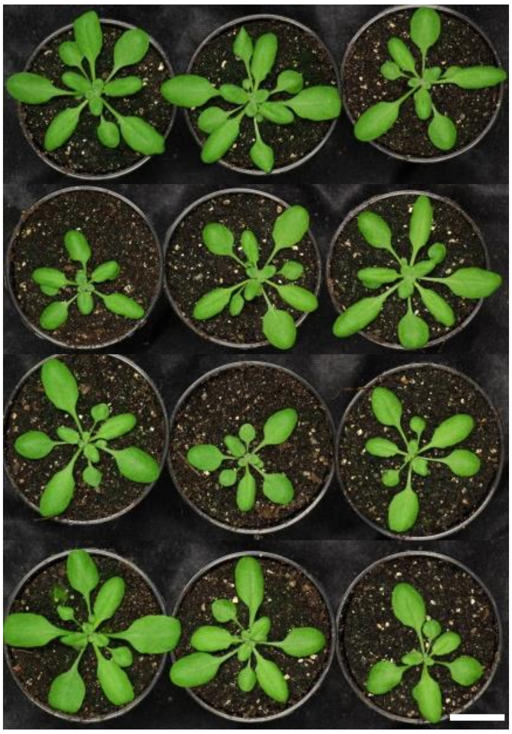

**Fig. S2** Rosette morphology of Arabidopsis plants expressing a heterologous TPP under the control of an axillary bud-specific promoter (long-day conditions). Supports Fig. 2.

The *Caenorhabditis elegans* TPP (CeTPP) were expressed in Arabidopsis Col-0 under the control of an axillary bud-specific promoter (*pBRC1*). Plants were grown in long-day conditions (16-h photoperiod), along with wild-type Arabidopsis Col-0 plants and a *pGLDPA:GUS* control line, and photographed at 22 DAS. Scale bar = 2 cm.

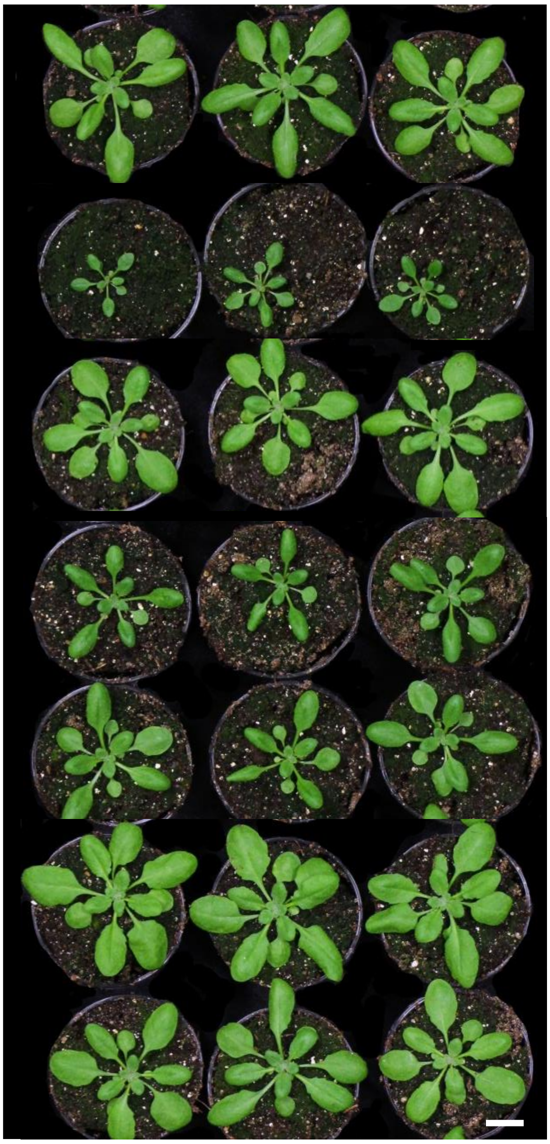

**Fig. S3** Rosette morphology of Arabidopsis plants expressing heterologous TPS or TPP under the control of constitutive or vasculature-specific promoters (long-day conditions). Supports Fig. 5.

The *Escherichia coli* TPS (OtsA) or the *Caenorhabditis elegans* TPP (CeTPP) were expressed in Arabidopsis Col-0 under the control of constitutive (*p35S*) or vasculature specific (*pGLDPA*) promoters. Plants were grown in long-day conditions (16-h photoperiod), along with wild-type Arabidopsis Col-0 plants and a *pGLDPA:GUS* control line. Rosette morphology at 22 DAS. Scale bar = 2 cm.

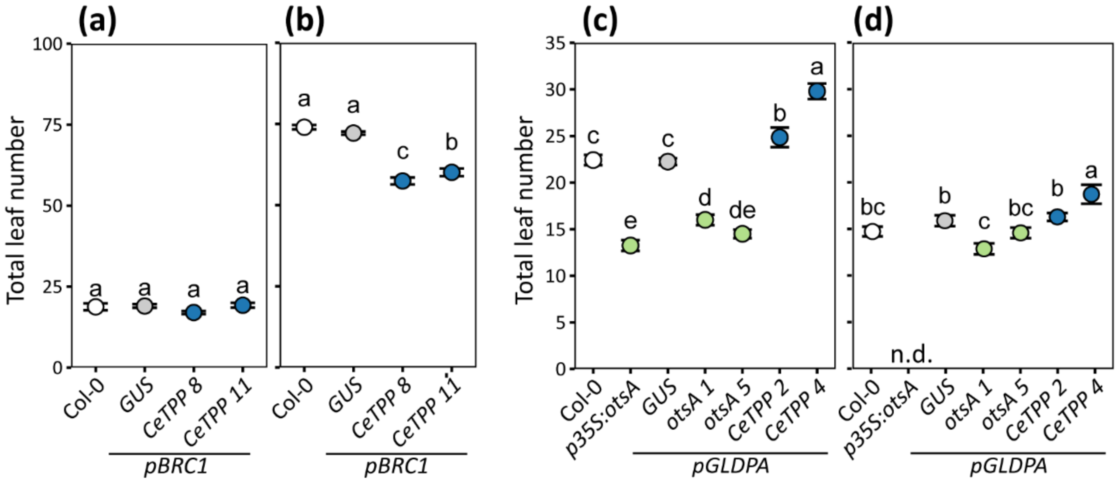

**Fig. S4** Flowering time of TPS and TPP over-expression lines. Supports Fig. 2, 5.

The $\beta$-*GLUCURONIDASE* (*GUS*), *Escherichia coli TPP* (*OtsA*) or *Caenorhabditis elegans TPP* (*CeTPP*) genes were expressed in Arabidopsis Col-0 under the control of an axillary bud-specific promoter (*pBRC1*) or a vasculature-specific promoter (*pGLDPA*) and flowering time was scored as total leaf number. The *pBRC1:CeTPP* and control plants were grown in (a) a 16-h or (b) 8-h photoperiod. The *pGLDPA:otsA/CeTPP* and control plants were grown in (c) a 16-h or (d) 18-h photoperiod. Wild-type and transgenic lines expressing heterologous proteins are represented by different symbol colours: Col-0 (white), GUS (grey), OtsA (green) and CeTPP (blue). Data are presented as mean ± S.E.M. ($n$ = 8-20), and letters indicate significant differences between genotypes according to one-way ANOVA with post hoc LSD testing ($p \leq 0.05$). n.d., not determined.

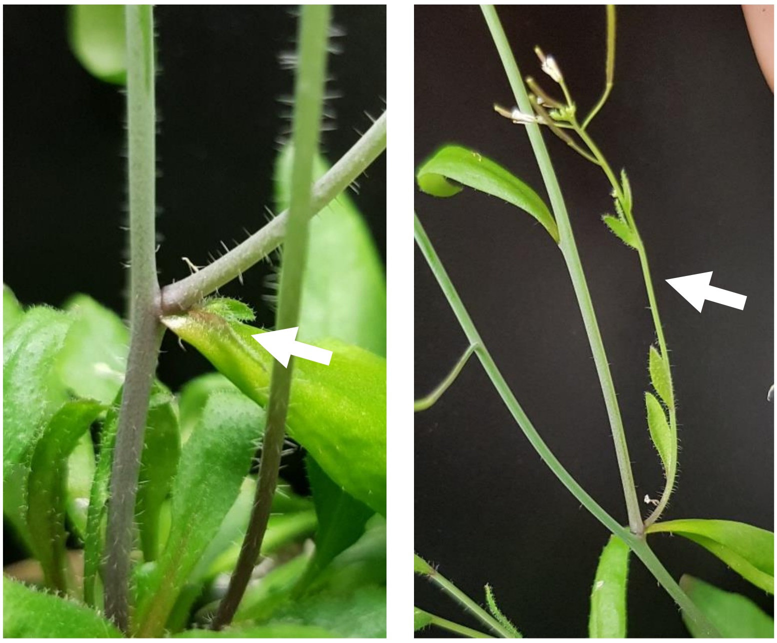

**Fig. S5** Cauline branching phenotype of Arabidopsis plants expressing a heterologous TPS under the control of constitutive or vasculature-specific promoters. Supports Fig. 5.

The *Escherichia coli* TPS (OtsA) was expressed in Arabidopsis Col-0 under the control of a vasculature specific (*pGLDPA*) promoters. Plants were grown in an 18-h photoperiod. Supports Fig. 5.Arrow point at second primary cauline branches.

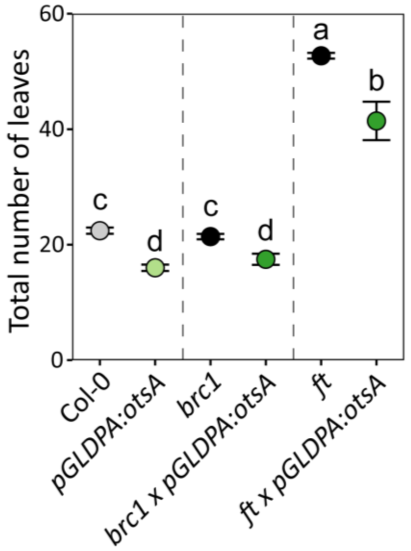

**Fig. S6** Effects of vasculature-specific TPS overexpression in wild-type and branching mutant backgrounds on flowering time under long-day conditions. Supports Fig. 7.

Arabidopsis *branched1* (*brc1*) and *flowering locus t* (*ft*) mutants and plants expressing the *Escherichia coli* TPS (OtsA) in the vasculature (*pGLDPA:otsA*) in these mutant backgrounds were grown in a 16-h photoperiod and flowering time recorded as total leaf number. Symbol colours: wild-type Col-0 (grey); *brc1* and *ft* parental mutants (black); *pGLDPA:otsA* expression in a wild-type background (light green); *pGLDPA:otsA* expression in a mutant background (dark green). Data are presented as mean ± S.E.M. ($n$ = 14-19) and letters indicate significant differences between genotypes according to one-way ANOVA with post hoc LSD testing ($p \leq 0.05$).

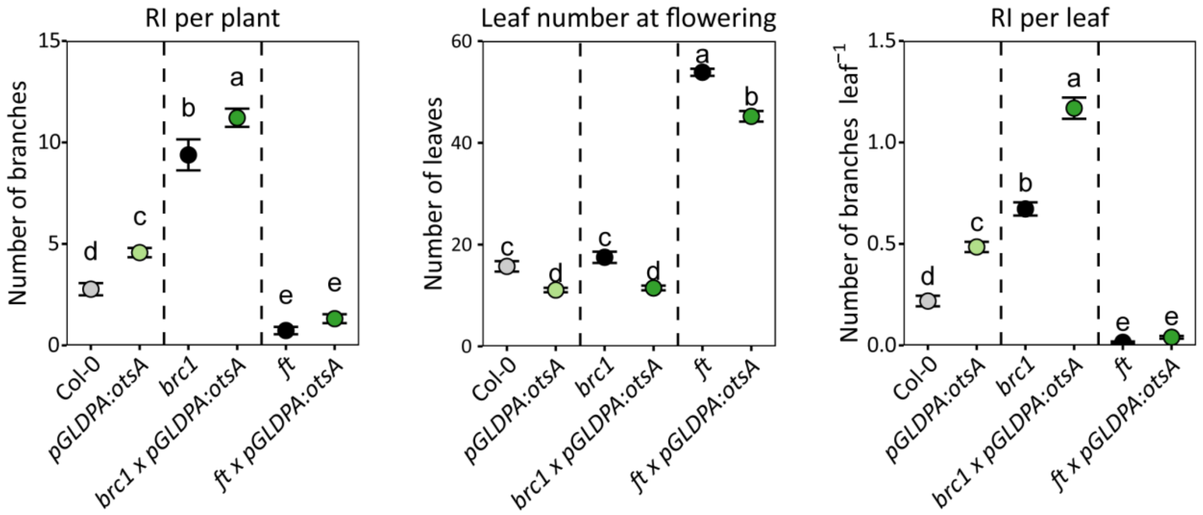

**Fig. S7** Effects of vasculature-specific TPS overexpression in wild-type and branching mutant backgrounds on flowering and shoot branching under long-day conditions (independent experiment supporting Fig. 7).

Primary rosette branches (RI) per plant, flowering time based on total leaf number and RI branches per rosette leaf in *branched1* (*brc1*) and *flowering locus t* (*ft*) mutants and plants expressing the *Escherichia coli* TPS (OtsA) in the vasculature (*pGLDPA:otsA*) in these mutant backgrounds. Symbol colours: wild-type Col-0 (grey); *brc1*, *max4* and *ft* mutants (black); *pGLDPA:otsA* expression in a wild-type background (light green); *pGLDPA:otsA* expression in a mutant background (dark green). RI branches (length ≥0.5 cm) were counted at the end of the plant's life cycle and normalized on rosette leaf number. Data are presented as mean ± S.E.M. ($n$ = 13-20) and letters indicate significant differences between genotypes according to one-way ANOVA with post hoc LSD testing ($p \leq 0.05$).

**Table S1** Oligonucleotide primers used for qRT-PCR analysis.

| Gene | Forward primer (5´→ 3´) | Reverse Primer (5´→ 3´) |
|---|---|---|
| **(i) Arabidopsis** | | |
| *ACTIN2*[a] | AGTGGTCGTACAACCGGTATTGT | GATGGCATGAGGAAGAGAGAAAC |
| *ACTIN7*[a] | AGTGGTCGTACAACCGGTATTGT | GAGGAAGAGCATACCCCTCGTA |
| *ACTIN8*[a] | AGTGGTCGTACAACCGGTATTGT | GAGGATAGCATGTGGAAGTGAGAA |
| *CO* | TCCATTAACCATAACGCATACATTT | CGGCACAACACCAGTTTCC |
| *FT* | ACCCTGGTGCATACACTGTT | GGTGGAGAAGACCTCAGGAA |
| *SUT1/SUC2* | GACGAACTATTCGGTGGTGGAA | CGCAATCGCTCCTAACACAAA |
| *SWEET11* | GACAAACCCCTAAACATGGCAA | TTCATGTAGCTGCTGCGGAAG |
| *SWEET12* | CAGAAGTGAGCATCGATATGGTGA | GCTACTGGTTCAGGAGATGTGAGTG |
| *SWEET13* | CTTCTACGTTGCCCTTCCAAATG | CTTTGTTTCTGGACATCCTTGTTGA |
| *SWEET14* | ACTTCTACGTTGCGCTTCCAAATA | CAGTTCAACATTAAAGTCAATCACTAATTC |
| *TSF* | GAAATGCCTTTGGCAATGAGGT | CCGGAACAATACCAACACAATACG |
| *TUBULIN3* | TGGTGGAGCCTTACAACGCTACTT | TTCACAGCAAGCTTACGGAGGTCA |
| **(ii) *E. coli*** | | |
| *otsA* | CGTTTTCTCGCCTATGAAGC | CCACGTCAGAGTAGCGGAAT |
| **(iii) Artificial** | | |
| *Spike 2* | ATTGCGCTCGCCATATACAC | GCTGGGATCAGGAGGAGAAG |
| *Spike 3* | GATCGTTTGCCTGCATTACC | GAGAGCGTCAGCCATACCAC |
| *Spike 4* | TACGTCGCACAACCACAATC | CAGCGCCACTAACCCACTAC |
| *Spike 5* | ACAAAGGCAGCGTTGAAAAC | TAGTGTCTGCACGCCATACC |
| *Spike 6* | CGCAAAGTCTCTCCTCTTGG | CAGTAGCCATTGCGGAAGAT |
| *Spike 7* | CTGAACCAGACTGCACATGG | ACGCTCATGGGCTTGTTTAT |

a alternative primer combinations used in the same reaction
